# Crosstalk between the Methyl-Cytosine Dioxygenase TET3 and the Methyl-CpG-binding protein MECP2 Controls Neuronal Maturation

**DOI:** 10.1101/2025.10.29.685266

**Authors:** Franziska R. Traube, Gilles Gasparoni, Anna Winkler, Anna S. Geserich, Hugo Sepulveda, J. Carlos Angel, Xiaojing Yue, Rouhollah Habibey, Victoria Splith, Gülce I. Gökҫe, Grazia Giorgio, Chiara Bernardini, Ruben Sachsse, Constanze Scheel, Marilla Bickerstaff-Westbrook, Martin Biel, Thomas Carell, Volker Busskamp, Anjana Rao, Jörn Walter, Stylianos Michalakis

## Abstract

Active DNA demethylation depends on Ten-Eleven-Translocation (TET) enzymes, which oxidize 5-methylcytosine (mC) to 5-hydroxymethylcytosine (hmC) and further derivatives. Mutations in *TET3*, encoding the predominant neuronal isoform, lead to Beck-Fahrner syndrome, a neurodevelopmental disorder. Using human iPSC-derived neurons, we show that TET3 is dispensable for neuronal specification but critical for subsequent maturation. Differentiating *TET3*-deficient neurons exhibit delayed transcriptional and proteomic transitions, altered synaptic signatures, and impaired network activity, indicating delayed functional maturation. Mechanistically, we identified an interaction between TET3 and the mC/hmC-binding protein MECP2, pathogenic variants of which cause Rett syndrome. MECP2 negatively regulates TET3 activity, as demonstrated in functional assays and by inverse hmC patterns in *MECP2*- and *TET3*-deficient neurons. Despite this, *MECP2*- and *TET3*-deficient neurons exhibit highly similar phenotypes later in differentiation. Our findings uncover a functional interplay between TET3 and MECP2 that coordinates DNA methylation and chromatin dynamics during neuronal maturation, suggesting a shared pathogenic mechanism in Beck-Fahrner and Rett syndromes.

## Introduction

Dynamic DNA methylation at cytosine (C) residues in the genome is a fundamental epigenetic mechanism in mammals, enabling cell type-specific gene expression patterns^1–4^. DNA methylation is established by a family of S-adenosyl-L-methionine-dependent DNA methyltransferases (DNMTs), which in humans includes three canonical members: DNMT1, DNMT3A, and DNMT3B. DNMT3A and DNMT3B are classified as *de novo* methyltransferases that introduce 5-methylcytosine (mC) at previously unmethylated CpG sites, while DNMT1 primarily maintains DNA methylation patterns by methylating hemi-methylated DNA following replication^5–7^. In mammalian genomes, mC is the most abundant DNA modification, comprising approximately 4 – 5% of all cytosines when normalized to genomic guanosine (G) content^8^. In regulatory and repetitive regions, mC is typically associated with gene silencing and correlates with repressive histone marks such as H3K9 methylation. In contrast, elevated DNA methylation is also found within the gene bodies of actively transcribed genes^8, 9^.

Active DNA demethylation is catalyzed by ten-eleven translocation (TET) enzymes, which oxidize mC to 5-hydroxymethylcytosine (hmC), and subsequently to 5-formylcytosine (fC) and 5-carboxycytosine (caC), in an α-ketoglutarate- and oxygen-dependent manner^10, 11^. The oxidized bases fC and caC can be excised and replaced by unmodified cytosine through the base excision repair pathway, specifically via thymine DNA glycosylase (TDG)^12^. In addition to its role as an intermediate in active DNA demethylation, hmC is also recognized as a stable epigenetic mark with distinct regulatory functions. In neurons, hmC is particularly abundant, up to ∼1% of cytosines, and is enriched at genes involved in synaptic function^13, 14^. Within CpG islands, hmC is associated with transcriptionally active chromatin and correlates positively with permissive histone modifications and negatively with repressive ones^15^. Brain-wide, hmC levels increase during neurogenesis and synaptogenesis, reaching their peak in the adult brain^16–18^. All three TET family members (TET1, TET2, and TET3), are expressed in the brain, but TET3 appears to play a particularly critical role during neurodevelopment ^19^. TET3 is the most highly expressed TET family member in neurons, and its haploinsufficiency in mice results in neonatal lethality^20^. In humans, heterozygous loss-of-function variants in *TET3* cause Beck-Fahrner syndrome (BEFAHRS), a neurodevelopmental disorder characterized by a spectrum of symptoms including developmental delay, intellectual disability, hypotonia, motor abnormalities, epilepsy, and ophthalmologic or growth impairments. Most pathogenic variants identified in BEFAHRS patients affect the catalytic domain or adjacent sequences of *TET3*^21^. Despite the significant role of TET3 in early brain development, simultaneous deletion of all three TET enzymes during the terminal differentiation of mouse Purkinje neurons reportedly causes only modest functional changes and transcriptional dysregulation in a subset of cell type-specific genes^22^. This observation suggests that TET3 function may be particularly crucial during earlier stages of neuronal differentiation.

To investigate the role of human TET3 during neuronal differentiation and maturation, we generated *TET3*-deficient human induced pluripotent stem cells (iPSCs) carrying a doxycycline-inducible *Neurogenin 1/2* expression cassette, which enables controlled neuronal reprogramming. This *Neurogenin*-induced neuronal differentiation system (hereafter abbreviated as iNGN) yields a highly homogeneous neuronal population within a few days of induction^23^.

In this study, we employ the iNGN system to demonstrate that *TET3* deficiency leads to delayed neuronal maturation. To identify proteins that may interact with TET3 and modulate its function, we further performed a screen for TET3-interacting partners at an early stage of neuronal maturation in the iNGN model. This screen revealed a previously uncharacterized interaction with methyl-CpG-binding protein 2 (MECP2), a key reader protein of both mC and hmC. Notably, MECP2 dysfunction causes Rett syndrome (RTT), a severe neurodevelopmental disorder^24–26^. We provide evidence for a functional interaction between TET3 and MECP2, which may contribute to the overlapping phenotypic manifestations observed in *TET3*- and *MECP2*-deficient states.

## Results

### The levels of hmC steadily increase during neuronal differentiation and maturation

Upon doxycycline induction of *Neurogenin 1* and *2*, iNGNs rapidly undergo neuronal reprogramming (Fig. S1A). As supported by previous literature^23^ and our immunofluorescence staining for proliferation and neuronal markers, the time points day 0 (d0), day 4 (d4), and day 8 (d8) represent key stages in the differentiation and maturation of iNGNs. In the uninduced state (d0), iNGN cells express proliferation markers, including Ki67, and incorporate 5-bromo-2’-deoxyuridine (BrdU), indicative of their proliferative, pluripotent status (Fig. 1A, d0). At d4 after doxycycline addition, iNGNs exhibit a characteristic neuronal morphology with neurite extensions that form a dense network and express the neuron-specific tubulin isoform beta 3 (TUBB3) (Fig. 1A, d4). At this intermediate stage, a subset of cells still expresses Ki67 and incorporates BrdU, indicating a transition phase during which cells are exiting the cell cycle and acquiring a postmitotic neuronal identity. By d8, expression of proliferation markers is no longer detectable, and the neuronal network appears more complex (Fig. 1A, d8). Although iNGNs do not yet exhibit action potentials at this stage^27^, morphological and molecular features suggest advanced neuronal maturation. Given the anticipated role of TET3 in early neuronal differentiation, we focused our subsequent analyses on these three developmental stages.

**Figure 1:**
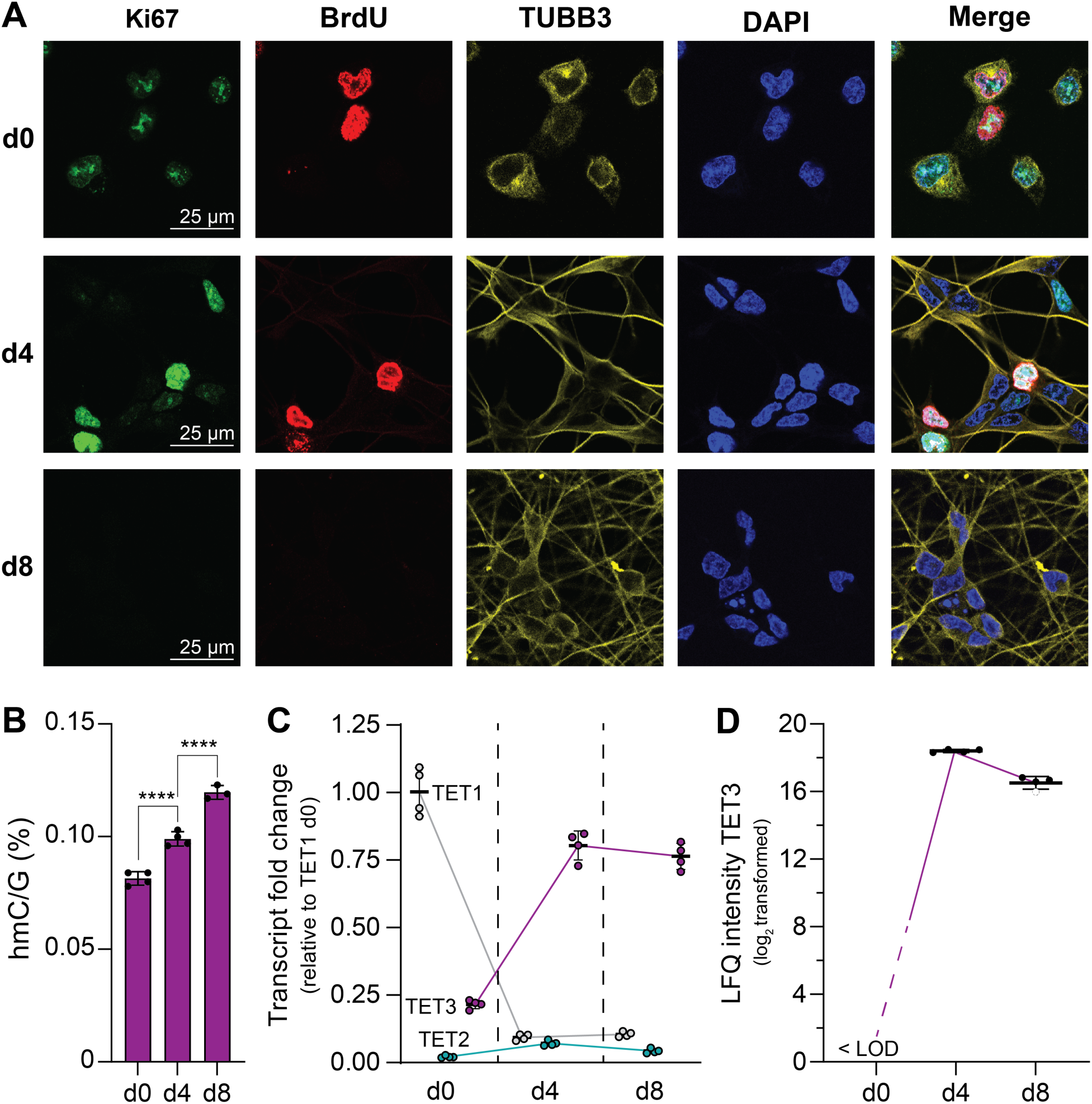
The iNGN model system to investigate the role of TET enzymes during neuronal differentiation and maturation. A) – D) iNGNs were analyzed at d0, d4 and d8 after induction of neuronal differentiation. A) Immunofluorescence staining against the proliferation markers Ki67 and BrdU as well as the neuron-specific marker TUBB3. DAPI was used to stain nuclei. One representative out of three biologically independent replicates is displayed. B) Levels of hmC normalized to G quantified by QQQ-MS. Bars represent mean. Ordinary one-way ANOVA combined with Šídák’s multiple comparisons test was performed. ****: p_adj_ < 0.0001, details are given in Supplementary Table 1. C) Transcript levels of *TET1*, *TET2* and *TET3* determined by RT-qPCR normalized to d0 *TET1* levels. D) Protein levels (log_2_ transformed LFQ intensities) of TET3 quantified by LC-MS/MS (LFQ-DIA). <LOD: below the limit of detection. Dashed circle at d8 indicates imputed log_2_ LFQ value. In B) – D), dots indicate data from biologically independent replicates. Error bars show S.D.

We found that the progression from pluripotent iPSCs (d0) to postmitotic neurons (d8) was accompanied by a steady increase in global genomic hmC levels quantified by triple-quadrupole mass spectrometry (QQQ-MS)^28^ (Fig. 1B). To determine whether this increase could be attributed to a specific TET enzyme, we quantified the transcript levels of *TET1*, *TET2*, and *TET3* at d0, d4, and d8 using quantitative real-time PCR (qRT-PCR) (Fig. 1C). *TET1* expression was high at d0 but declined sharply after induction, while *TET2* remained low throughout all time points. In contrast, *TET3* expression increased substantially from d0 to d4 and remained elevated through d8. These findings suggest that *TET3* is the dominant TET enzyme during early neuronal differentiation in this model.

This dynamic regulation of *TET3* was further confirmed at the protein level by liquid chromatography–tandem mass spectrometry (LC-MS/MS), which revealed undetectable TET3 protein at d0 and a substantial expression following neuronal reprogramming (Fig. 1D).

These observations indicate that TET3 is the primary driver of hmC accumulation during early neuronal differentiation and could potentially act as a key regulator of epigenetic reprogramming during early neuronal maturation.

### *TET3* deficiency delays neuronal maturation

To investigate the role of TET3 during early neuronal differentiation and maturation, we generated a *TET3* knockout (TET3^KO^) in the iNGN model by introducing a frameshift mutation within the catalytic domain of the *TET3* gene (Fig. S1B) and confirmed absence of wild-type (WT) *TET3* transcript (Fig. S1C) and absence of TET3 on the protein level (Fig. S1D). We first assessed the impact of TET3 loss on neuronal differentiation using brightfield microscopy (Fig. 2A). Morphological assessment revealed no overt differences between WT and TET3^KO^ iNGNs during the early stages of differentiation. TET3^KO^-derived neurons extended axons and formed neuronal networks by d4, similar to WT cells (Fig. 2A). However, despite equivalent seeding densities at d0, TET3^KO^ cultures appeared to contain fewer cells by d4 and d8, as observed in brightfield microscopy. This impression was confirmed by independent cell counting, which showed a significantly reduced number of cells in TET3^KO^ cultures following differentiation (Fig. S1E).

**Figure 2:**
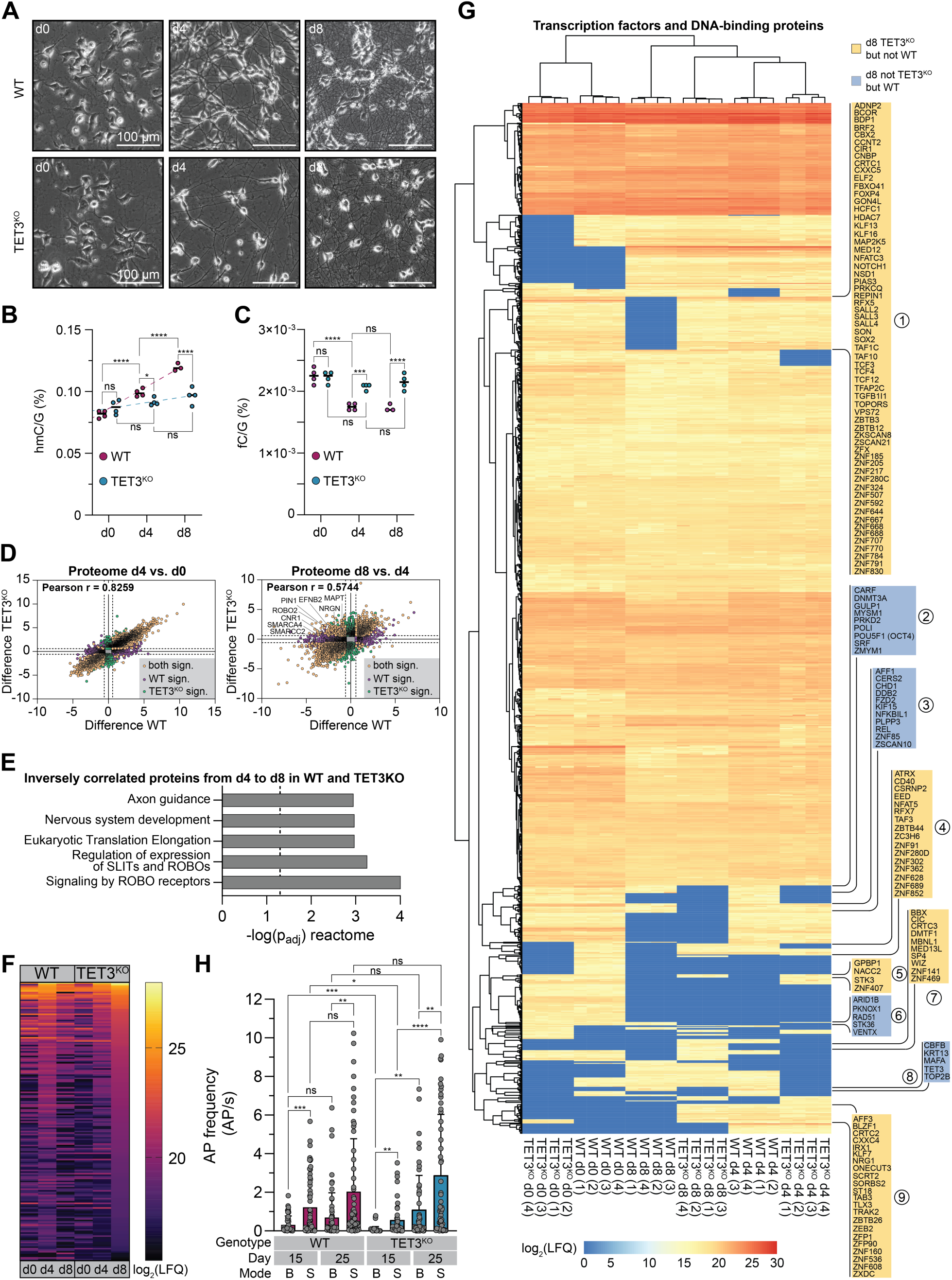
Differentiation and maturation of TET3^KO^ iNGNs compared to WT. A) Brightfield microscopy images of WT and TET3^KO^ iNGNs at d0, d4, and d8. One out of three biologically independent replicates is displayed. B), C) Quantification of genomic hmC (B) and fC (C) levels, normalized to G by QQQ-MS in WT and TET3^KO^ iNGNs at d0, d4, and d8. Dots represent biologically independent replicates. Statistical analysis was performed using ordinary two-way ANOVA followed by Tukey’s multiple comparisons test (between genotypes at each time point and across time points within genotypes). Details are given in Supplementary Table 1. Data for hmC in WT are also displayed in Fig. 1B. D) – G) Quantitative proteomic analysis using LFQ-DIA mass spectrometry (four biological replicates per genotype and time point). D) Correlation plots of proteomic changes from d0 to d4 (left) and d4 to d8 (right) in WT and TET3^KO^ iNGNs. Four biologically independent replicates were analyzed per genotype per timepoint. Significantly regulated proteins were identified using a permutation-based t-test with thresholds of –log_10_(p-value) > 1.3 and difference (|log_2_FC|) > 0.58496 (fold change > 1.5). E) Pathway enrichment analysis of proteins showing inverse expression changes from d4 to d8 in WT vs. TET3^KO^, using the Reactome pathway knowledgebase^48^. F) Heatmap of mean LFQ values of these inversely regulated proteins at d0, d4, and d8 in WT and TET3^KO^ iNGNs. G) Hierarchical clustering (Euclidean distance) based on transcription factors and DNA-binding proteins identified in the proteomic dataset. H) Action potential (AP) frequency under baseline (B) and optogenetically stimulated (S) conditions at d15 and d25. Data were obtained from four independent MEA cultures for WT and from three for TET3^KO^. Brown-Forsythe and Welch ANOVA tests with Dunnett’s T3 multiple comparisons test was performed. B), C), H) p_adj_: ns ≥ 0.05, * < 0.05, ** < 0.01, *** < 0.001, **** < 0.0001 and details are given in Supplementary Table 1.

We next quantified the levels of hmC and found that TET3^KO^ cells showed a significantly smaller increase in hmC compared to WT cells during differentiation (Fig. 2B). At d0, hmC levels were slightly elevated in TET3^KO^ cells compared to WT, but this difference was not statistically significant. However, by d4, WT cells had significantly higher hmC levels, and this difference became even more pronounced by d8. In contrast, global fC levels displayed an inverse trend: both WT and TET3^KO^ cells started with comparable levels at d0, but fC levels declined significantly in WT cells over time, while remaining relatively stable in TET3^KO^ cells (Fig. 2C). These data suggest that TET3 is the principal TET enzyme responsible for the dynamic regulation of hmC and fC during neuronal differentiation and maturation.

To gain a broader perspective on the molecular consequences of TET3 loss, we performed whole-cell proteome profiling at d0, d4, and d8 in both WT and TET3^KO^ iNGNs using label-free quantification (LFQ). We compared protein expression changes over two key intervals – d0 to d4 (differentiation phase) and d4 to d8 (early maturation phase) – between WT and TET3^KO^ cells (Fig. 2D, Supplementary Table 2). Proteomic changes during the initial phase (d0 to d4) were highly correlated between genotypes, indicating that *TET3* deficiency has only a modest impact on the early stages of neuronal differentiation. In contrast, the transition from d4 to d8 revealed substantial divergence between WT and TET3^KO^ cells. Notably, a subset of proteins was significantly downregulated in WT cells from d4 to d8 but upregulated in TET3^KO^ cells over the same interval (Fig. 2D, Supplementary Table 2).

Pathway enrichment analysis of these inversely regulated proteins identified significant overrepresentation of Gene Ontology (GO) terms related to axonal development and nervous system maturation (Fig. 2E, Supplementary Table 2), suggesting that *TET3* deficiency disrupts molecular programs essential for neuronal network formation. Among the proteins that were significantly downregulated in WT iNGNs from d4 to d8, but upregulated in TET3^KO^ iNGNs were several proteins involved in axonal cytoskeletal stabilization and guidance, such as the microtubule-associated proteins tau (MAPT) as well as the tau-regulating prolyl isomerase PIN1^29,30^. Also included were several proteins implicated in axon guidance, synapse density and plasticity, such as ephrin-B2 (EFNB2) and neurogranin (NRGN), receptor proteins with key roles in neuronal signaling, such as roundabout guidance receptor 2 (ROBO2) and cannabinoid receptor 1 (CNR1), as well as chromatin remodeling proteins essential for neuronal differentiation and development, including components of the BAF (SWI/SNF) complex, SMARCA4 and SMARCC2^31, 32^.

To gain a more comprehensive understanding of how proteins exhibiting inverse expression changes between WT and TET3^KO^ iNGNs from d4 to d8 differ in their expression dynamics over the full course of differentiation and maturation (d0 to d8), we ranked these proteins based on their LFQ intensities on d8 in TET3^KO^ cells and analyzed their expression patterns at d0, d4, and d8 in both genotypes (Fig. 2F). On d0, LFQ values for these proteins were generally low in both WT and TET3^KO^ cells, indicating no or minimal expression in the pluripotent stem cell state. Of note, this subset of proteins showed an increase in abundance from d0 to d4 in WT cells, followed by a decline from d4 to d8. In TET3^KO^ cells, their expression also increased from d0 to d4, but to a lesser extent. However, their expression levels in TET3^KO^ iNGNs further increased from d4 to d8 and reached levels comparable to those observed in WT cells at d4 (Fig. 2F). Thus, proteins implicated in early neuronal maturation in WT iNGNs, showed a delayed increase in abundance in TET3^KO^ iNGNs, suggesting a delay or impairment of neuronal maturation in these cells.

We next extended the analysis to all transcription factors and DNA-binding proteins^33^ robustly detected in at least one genotype, defined as presence in ≥3 of 4 biologically independent replicates at any time point. To distinguish low-level expression from likely absence, i.e., undetected in all replicates, we applied an imputation strategy assigning a log2-transformed LFQ value of 1 to the latter group. This allowed proper representation of true absence in downstream analyses. Hierarchical clustering using Euclidean distance confirmed high intra-group consistency (Fig. 2G), with replicates clustering by genotype and time point. As expected, d0 samples from both WT and TET3^KO^ iNGNs clustered together, reflecting their undifferentiated state. In contrast, d4 and d8 samples formed a distinct, neuronal cluster. Within this cluster, d4 WT and TET3^KO^ samples grouped together, while d8 TET3^KO^ samples clustered closer to d4 WT and TET3^KO^ samples than to d8 WT samples, which formed a separate subcluster. This pattern confirmed a delayed maturation trajectory in TET3^KO^ iNGNs, with their transcription factor and DNA-binding protein profiles at d8 remaining more similar to d4 than to fully differentiated WT neurons.

To further characterize genotype-specific transcription factor and DNA-binding protein profiles, we analyzed proteins detected at d8 in only one genotype. Based on their temporal expression patterns, we identified nine distinct clusters (Supplementary Table 2).

Cluster 1, the largest group, comprised transcription factors and DNA-binding proteins non-detectable at d8 in WT cells but present at all other time points in both genotypes. This cluster included transcription factors characteristic of pluripotent or progenitor states and important for neurodevelopment, such as SOX2, which maintains neural stem cell identity and regulates the transition from proliferation to differentiation^34^, and NOTCH1, a key mediator of neuronal fate decisions^35^. SALL4, a zinc finger transcription factor implicated in neural progenitor maintenance and reported to co-bind hmC and TET enzymes^36^, was also included. Cluster 2 contained proteins expressed at d0 in both genotypes but undetectable in TET3^KO^ iNGNs at d4 and d8, while remaining present in WT throughout differentiation. Notably, this cluster included DNMT3A, a DNA methyltransferase highly expressed in postmitotic neurons and involved in transcriptional regulation and synaptic plasticity^37–40^. Its expression normally increases during neuronal differentiation, correlating with higher mC levels in neurodevelopmental genes^37, 41, 42^. POU5F1 (OCT4), a core pluripotency factor ^43^, also remained expressed in WT but not TET3^KO^ iNGNs during differentiation. Cluster 3 comprised proteins absent at d8 in TET3^KO^ cells but present at all other time points in both genotypes. An example is the Frizzled receptor FZD2, a Wnt receptor involved in neuronal differentiation, migration, and synaptogenesis^44^. Cluster 4 included proteins with more complex expression dynamics that were present in WT at d4 and absent at d8, while they were undetectable in TET3^KO^ iNGNs at d4, but detectable at d8. Among them was ATRX, a chromatin remodeler implicated in neurogenesis, synaptic development, and neuronal survival, and known to interact with SOX2 during neuronal differentiation ^45^. Clusters 5 and 6 contained few proteins and cluster 7 comprised transcription factors and DNA-binding proteins exclusively detected at d8 in TET3^KO^ cells. Among them was SP4, a neuron-specific transcription factor involved in regulating genes associated with synaptic plasticity, dendritic morphology, and ion channel expression^46^. Cluster 8 included proteins absent in TET3^KO^ iNGNs at all time points but present in WT from d4 onward. This group includes TET3 itself, consistent with its knockout status. Cluster 9 contained proteins undetectable at d0 in both genotypes, expressed at d4, but subsequently lost in WT by d8 while persisting in TET3^KO^. An example is KLF7, a zinc finger transcription factor essential for axon guidance and neurite outgrowth^47^.

Previous studies have shown that iNGN-derived neurons co-cultured with astrocytes are capable of firing action potentials by d14, and that more mature neurons exhibit spontaneous postsynaptic currents indicative of functional synaptic activity. To determine whether the delayed maturation observed in TET3^KO^ iNGNs at the proteomic level is reflected in neuronal network function, we assessed spontaneous (baseline) and optogenetically evoked (stimulated) activity using microelectrode array (MEA) measurements. To this end, iNGNs were transduced on d1 with a lentiviral construct encoding EYFP-tagged ChRimson for optical stimulation. On d5, cells were re-seeded onto MEA chips in co-culture with astrocytes and maintained in medium supporting neuronal network activity (Fig. S2A – D). Spontaneous and optogenetically-evoked activity was subsequently recorded on d15 and d25 in both WT and TET3^KO^ iNGNs (Fig. 2H). In both genotypes, optogenetic stimulation significantly increased the action potential (AP) frequency at both time points compared to baseline. However, spontaneous and evoked activity were significantly lower in TET3^KO^ neurons compared to WT on d15. From d15 to 25, WT iNGNs showed only a modest, non-significant increase in AP frequency, whereas TET3^KO^ iNGNs exhibited a significant increase in both spontaneous and evoked activity, reaching the corresponding WT levels by d25 (Fig. 2H). These findings indicate that the delayed maturation of TET3^KO^ iNGNs is associated with a corresponding delay in the functional establishment of neuronal network activity.

### MECP2 is an interaction partner of TET3 that modulates TET3 catalytic activity

Proteins function within complex interaction networks rather than in isolation. To characterize the TET3 interactome during neuronal maturation, we performed co-immunoprecipitation (co-IP) experiments in d8 WT iNGNs using two complementary approaches: GFP-TET3-based coIP with enrichment of TET3 fusion protein (Fig. 3A)^49^, and coIP of endogenous TET3 (Fig. 3B, Fig. S3A). In the GFP-TET3 coIP, interaction partners were identified by LC-MS/MS analysis followed by statistical enrichment analysis using a t-test with permutation-based FDR calculation using GFP control coIP as the reference (Fig. 3A). Among the enriched proteins were several chromatin-associated factors, including all three components of the FET complex - FUS, EWS, and TAF15 - which are known to link chromatin-mediated transcriptional regulation to RNA processing and have established roles in neuronal differentiation and maturation^50^. Also highly enriched were high mobility group box proteins HMGB1 – 3, which participate in chromatin remodeling and transcriptional co-activation and are implicated in neuronal development and neurite outgrowth^51^. Additionally, we identified the chromobox proteins CBX1, CBX3, and CBX5, which recognize H3K9me2/3 histone marks and coordinate key chromatin regulatory processes during neurodevelopment^52^. Most notably, we identified MECP2 as a significantly enriched interaction partner of TET3 in the GFP-TET3 coIP (Fig. 3A). MECP2 is a known reader of mC and hmC with transcriptional regulatory functions and critical roles in neuronal development and function^53, 54^. Loss-of-function mutations in the X-linked *MECP2* gene cause RTT, a severe neuronal developmental disorder affecting approximately 1 in 10,000 female births^24–26^. RTT patients typically exhibit apparently normal development during the first 6 –18 months of life, followed by progressive regression in acquired motor and language skills. After a transient period of stabilization, neurological deterioration continues^55^.

**Figure 3:**
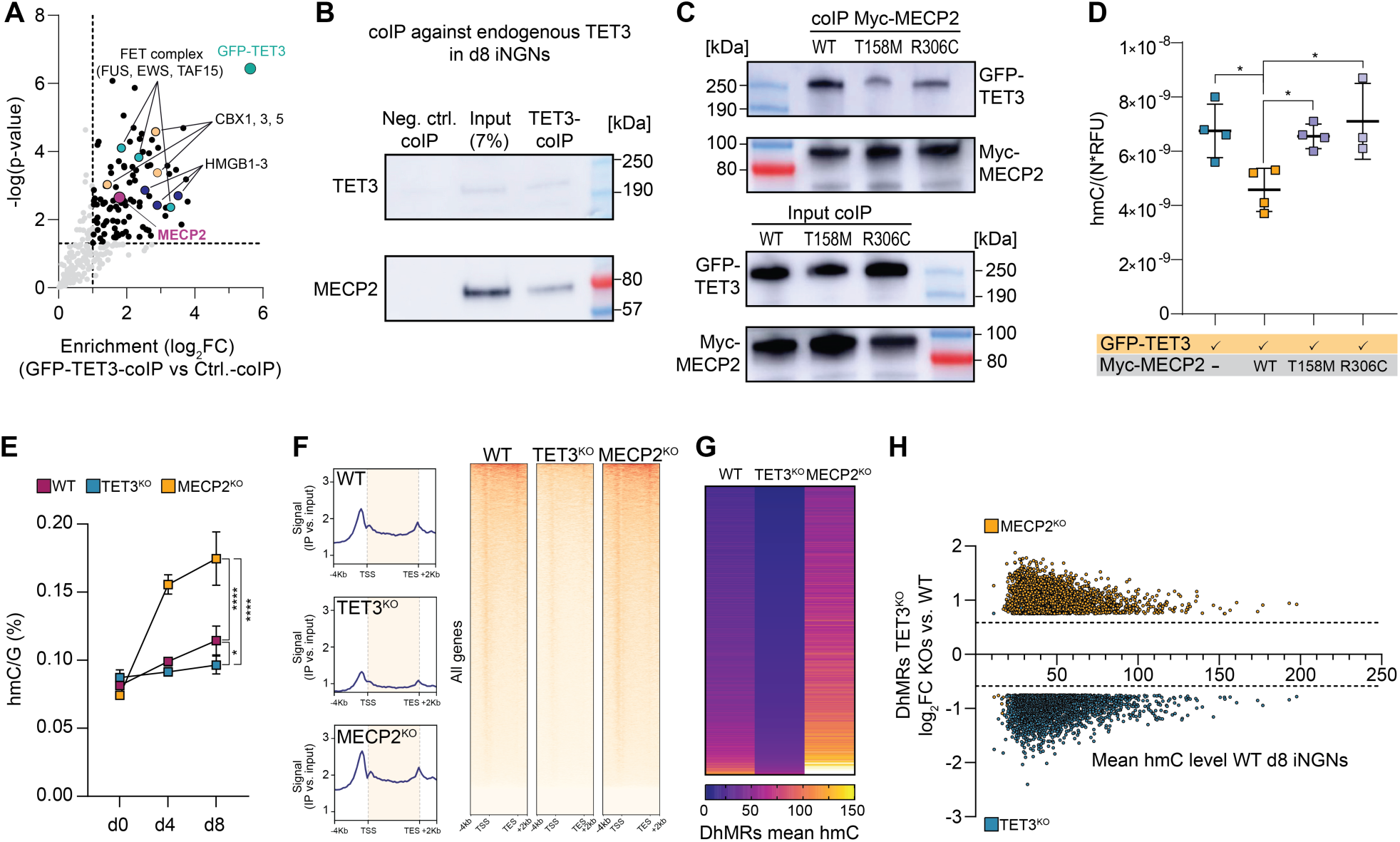
TET3–MECP2 interaction and its functional impact on hmC levels. A) GFP-TET3 coIP in nuclear lysates of d8 WT iNGNs, analyzed by MS. Four biologically independent samples were analyzed per sample type. Proteins significantly enriched relative to GFP control (t-test with permutation-based FDR calculation; significance thresholds – log(p-value) > 1.3 and log₂ fold change > 1 (fold change > 2.0)) are shown in black or highlighted in color. B) CoIP of endogenous TET3 from d8 WT iNGN nuclear lysates, analyzed by immunoblotting. IgG pulldown served as a negative control; 35 µg of nuclear lysate was used as input (positive control). One out of two independent experiments is displayed. C) HEK293T cells were co-transfected with plasmids encoding GFP-TET3 and either Myc-tagged WT MECP2 or two RTT syndrome-associated MECP2 mutants (T158M and R306C). After 48 h, Myc-MECP2 variants were immunoprecipitated, and co-precipitation of GFP-TET3 was assessed by immunoblotting alongside the input samples for coIP (4%). One out of three independent experiments is displayed. D) In parallel with the experiment in panel C, genomic DNA was isolated and global hmC levels were quantified by QQQ-MS. Values were normalized to canonical DNA nucleosides (per N) and GFP fluorescence intensity^49^, and each dot represents an independent biological sample. E) Quantification of global hmC levels at d0, d4, and d8 in WT, TET3^KO^, and MECP2^KO^ iNGNs (n = 4 for WT and TET3^KO^ and n = 3 for MECP2^KO^; dots represent mean, error bars represent S.D.). WT data from Fig. 1B and 2B and TET3^KO^ data from Fig. 2B are shown again here for direct comparison with MECP2^KO^. F) Heatmap visualization of hmC mapping in d8 iNGNs showing tracks for CMS-IPs subtracting input. For each genotype (n = 2), average levels were calculated. G), H) Average levels of the top 3978 common DhMRs in d8 TET3^KO^ vs. WT iNGNs and how these regions are hydroxymethylated in MECP2^KO^, displayed as heatmap (G) and scatter plot (H). D), E) p_adj_: ns ≥ 0.05, * < 0.05, **** < 0.0001. Ordinary one-way ANOVA combined with Tukey’s multiple comparisons test was performed and details are given in Supplementary Table 1.

To validate the interaction between TET3 and MECP2, we performed coIP with endogenous TET3 followed by immunoblotting, confirming MECP2 as a binding partner (Fig. 3B, Fig. S3A). We then conducted a reverse coIP in HEK293T cells co-expressing GFP-TET3 and Myc-tagged MECP2. Myc-IP was performed using either WT MECP2 or two RTT-associated MECP2 missense mutants: T158M, which impairs DNA binding, and R306C, which disrupts other known protein– protein interactions and transcriptional regulatory function^56^. While GFP-TET3 was strongly enriched in the immune-precipitate of MECP2^WT^, its enrichment was substantially reduced in the MECP2^T158M^ and MECP2^R306C^ pulldowns, despite comparable input levels of both GFP-TET3 and Myc-MECP2 (Fig. 3C, Fig. S3B). To explore the functional consequences of this interaction, we quantified genomic hmC levels by QQQ-MS in HEK293T cells co-expressing GFP-TET3 and either MECP2^WT^, MECP2^T158M^, or MECP2^R306C^, normalizing for GFP-TET3 expression according to the GFP signal. Co-expression with MECP2^WT^ reduced TET3 catalytic activity, as evidenced by significantly decreased hmC levels, whereas co-expression with either RTT-associated mutant had no such effect (Fig. 3D). These results suggest that MECP2 competes with TET3 at mC-enriched genomic loci, and that this competition requires intact DNA- and protein-binding capacity of MECP2.

To examine the role of MECP2 in regulating TET3 activity in a neuronal context, we generated an *MECP2* knockout iNGN line (MECP2^KO^; Fig. S3C, D) and quantified global hmC levels at d0, d4, and d8 of differentiation in WT, TET3^KO^, and MECP2^KO^ iNGNs by QQQ-MS (Fig. 3E, Fig. S3E). In MECP2^KO^ cells, neuronal induction was associated with a significantly greater accumulation of hmC compared to WT, consistent with increased TET activity in the absence of MECP2, as suggested by the HEK293T-based assay (Fig. 3D).

Finally, to assess site-specific changes in hmC, we performed chemical labeling of hmC through sodium bisulfite-mediated conversion to cytosine-5-methylenesulfonate (CMS), followed by CMS-IP and spike-in normalization^57, 58^ comparing d8 WT, TET3^KO^, and MECP2^KO^ iNGNs. Global analysis of CMS-IP results across all genes confirmed our result obtained by QQQ-MS that TET3^KO^ iNGNs featured reduced hmC levels at d8, while MECP2^KO^ iNGNs feature increased hmC levels (Fig. 3F). We identified 15,506 differentially hydroxymethylated regions (DhMRs) in TET3^KO^ compared to WT. Of these, 15,475 regions showed significantly reduced hmC levels, while only 31 regions were hyper-hydroxymethylated (Fig. S3F). DhMRs were broadly distributed across genomic contexts, including intronic sequences (35% of downregulated and 26% of upregulated DhMRs), intergenic regions (24% down, 16% up), long terminal repeats (LTRs; 9% down, 23% up), long interspersed nuclear elements (LINEs; 8% down, 19% up), short interspersed nuclear elements (SINEs; 8% down), and promoter regions (4% down) (Fig. S3G). In MECP2^KO^, we detected 43,419 DhMRs with increased hmC relative to WT and only 63 with decreased levels (Fig. S3H). These DhMRs showed a similar genomic distribution to those in the TET3^KO^ (Fig. S3G). Notably, 11,543 DhMRs were shared between the TET3^KO^ vs. WT and MECP2^KO^ vs. WT comparisons, representing 75% of all DhMRs in the TET3^KO^. These overlapping regions exhibited a substantial loss of hmC in the TET3^KO^ and a gain in hmC in the MECP2^KO^ (Fig. 3G, H, Fig. S3F). This inverse relationship indicates that TET3 and MECP2 co-occupied common genomic regions and functionally interact, in line with our proteomic findings. Moreover, the hmC sequencing analysis supports our QQQ-MS data, demonstrating that while *TET3* deletion leads to a global reduction of hmC, *MECP2* deficiency enhances TET enzyme activity.

### *TET3* and *MECP2* deficiencies result in highly similar phenotypes in post-mitotic immature neurons

We next examined how *MECP2* deficient neurons compare phenotypically with WT and TET3^KO^ neurons. Across the differentiation timeline from d0 to d8, the morphology of WT, TET3^KO^, and MECP2^KO^ cells appeared broadly similar, with no obvious differences in cell shape. However, by d8, not only TET3^KO^ but also MECP2^KO^ exhibited substantially less dense neuronal networks than WT, as assessed by brightfield microscopy (Fig. S4A). This observation was confirmed by quantification of cell numbers at d0, d4, and d8 (Fig. 4A). While WT iNGNs showed a significant increase in cell number following the induction of neuronal differentiation, TET3^KO^ cells exhibited only a modest, non-significant increase, while MECP2^KO^ cells showed no increase at all. By d8, both KOs displayed significantly reduced cell numbers compared to WT (Fig. 4A).

**Figure 4:**
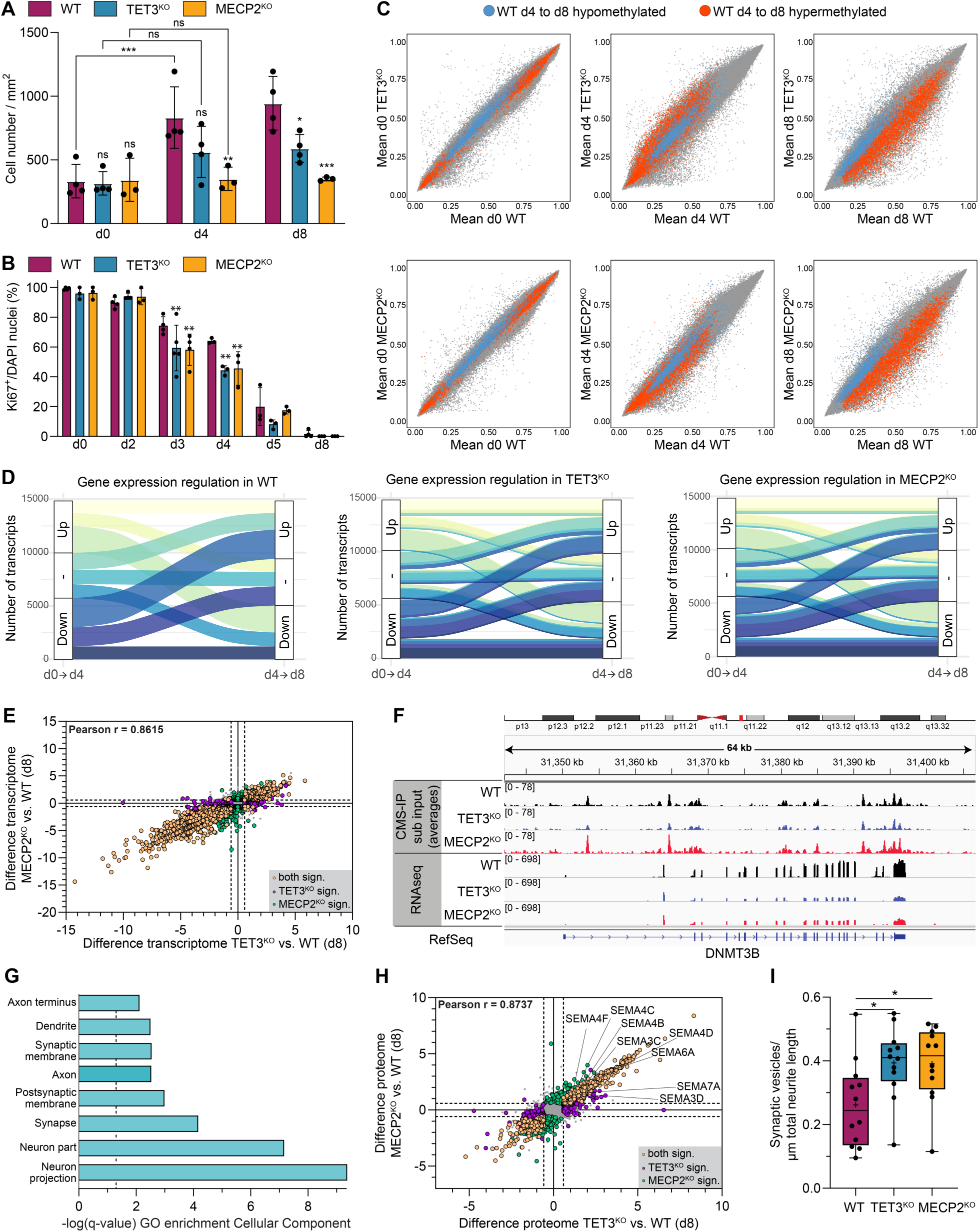
Phenotypic convergence of TET3^KO^ and MECP2^KO^ iNGNs by d8 of neuronal differentiation and comparison to WT. A) Cell counts at d0, d4 and d8. Data for WT and TET3^KO^ are also displayed in Fig. S1E. Bars represent the mean ± S.D.; each dot corresponds to a biologically independent replicate. Significance annotations above bars refer to comparisons with WT. Statistical analysis was performed using ordinary two-way ANOVA followed by Tukey’s multiple comparisons test (between genotypes at each time point and across time points within genotypes). Details are given in Supplementary Table 1. B) Proportion of Ki67-positive cells during early differentiation. Bars represent the mean ± S.D.; each dot represents a biologically independent replicate. Statistical analysis was performed using ordinary two-way ANOVA followed by Tukey’s multiple comparisons test (between genotypes at each time point). Details are given in Supplementary Table 1. C) DNA methylation profiles (EPIC array) shown as average methylation correlation plots across genotypes and time points (≥3 biologically independent replicates per group). D) Alluvial plots showing transcriptional transitions between d0–d4 and d4–d8 based on RNA-seq (3 biologically independent replicates per group). Genes were classified as upregulated (log₂FC > 0.58496, p_adj_ < 0.05), downregulated (log₂FC < –0.58496, p_adj_ < 0.05), or unchanged (“–”). Color assignments were defined for the WT and applied consistently across genotypes to enable direct comparison. E) Correlation plot of differential gene expression (log₂FC) between each KO and WT at d8. RNA-seq data from n = 3 biologically independent replicates per genotype. DEGs (p_adj_ < 0.05 and |log₂FC| > 0.58496) are highlighted as indicated. F) Genomic tracks showing hmC signal and RNA-seq coverage at the *DNMT3B* locus. G) GO cellular component enrichment analysis of genes upregulated in both TET3^KO^ and MECP2^KO^ compared to WT at d8. H) Correlation plot of differential protein expression (log₂FC) between each KO and WT at d8, based on global proteomic analysis (n = 4 biologically independent replicates per genotype). Differentially expressed proteins (permutation-based t-test, p < 0.05 and |log₂FC| > 0.58496) are highlighted in color. I) Quantification of synaptic vesicle density (SV2A-positive puncta per µm neurite length) in d8 iNGNs, assessed by immunofluorescence (SV2A and TUBB3 staining). Each dot represents an independent imaging field; data derived from three biologically independent samples per genotype. A), B), I) p_adj:_ ns > 0.05, * < 0.05, ** < 0.01, *** < 0.001, **** < 0.0001; details are given in Supplementary Table 1.

To determine whether the reduced cell numbers in TET3^KO^ and MECP2^KO^ iNGNs resulted from altered proliferation dynamics, we analyzed Ki67 expression (Fig. 4B) and BrdU incorporation (Fig. S4B). At d0, all genotypes showed high proliferation, with ∼60% of cells incorporating BrdU and universal Ki67 positivity. By d2, although all cells remained Ki67-positive, BrdU incorporation declined to ∼30% in WT, reflecting a slowdown in replication. Interestingly, MECP2^KO^ cells retained significantly higher BrdU incorporation (∼40%), suggesting delayed deceleration of proliferation in the early differentiation state. TET3^KO^ cells showed BrdU levels similar to WT at both d0 and d2. As differentiation progressed, BrdU incorporation declined in all genotypes, but more steeply in both KOs from d3 onward. By d4, BrdU and Ki67 levels were significantly lower in TET3^KO^ and MECP2^KO^ compared to WT, indicating earlier cell cycle exit. Despite this, early BrdU incorporation was still detectable, suggesting that reduced cell density in the KOs cannot be attributed solely to decreased proliferation. Instead, additional cell loss appears to contribute to the lower cell numbers observed during differentiation in both TET3^KO^ and MECP2^KO^ lines.

Overall, TET3^KO^ and MECP2^KO^ neurons exhibit surprisingly similar phenotypes during differentiation, including reduced cell proliferation and early cell cycle exit. However, these similarities occur in the context of opposing hmC profiles. To determine whether other molecular layers, namely DNA methylation, gene expression and proteome composition, could explain this shared phenotype, we systematically profiled all three across differentiation.

Genome-wide DNA methylation was assessed using the Infinium MethylationEPIC BeadChip, covering over 850,000 CpG sites (Supplementary Table 3). Principal component (PC) analysis (PCA) revealed a clear separation of undifferentiated iNGNs (d0) from differentiating cells (d4 and d8) along PC1 (Fig. S4C), corresponding to global methylation changes during lineage commitment. PC2 further separated d4 from d8 samples. At d0, all genotypes overlapped in the PCA space, consistent with low TET3 and MECP2 expression levels in pluripotent cells. By d4, WT, TET3^KO^, and MECP2^KO^ formed distinct clusters, indicating genotype-specific methylation profiles. At d8, WT samples shifted further along PC2, while TET3^KO^ and MECP2^KO^ remained closer to d4 and, notably, clustered together, suggesting a shared, delayed methylation trajectory in both KOs. At the single-site level, we observed high correlation of CpG methylation across all genotypes at d0 (Fig. S4D, Supplementary Table 3). By d4, CpGs were increasingly differentially methylated in TET3^KO^ and in MECP2^KO^ compared to WT, but also between the two KOs. These differences suggest initially distinct methylation programs. However, by d8, methylation changes relative to WT persisted, while the profiles of both KOs converged, showing near-identical methylation landscapes (Fig. S4D, Supplementary Table 3).

We next examined DNA methylation dynamics over time, specifically from d0 to d4 and from d4 to d8 to determine how methylation trajectories in TET3^KO^ and MECP2^KO^ compare to WT. During early differentiation (d0 to d4), the top 10,000 CpG sites undergoing demethylation in WT cells were not equivalently demethylated in TET3^KO^, which retained higher methylation levels at these sites on d4 (Fig. S4E, Supplementary Table 3). However, by d8, methylation at these sites in TET3^KO^ had largely reached WT levels, consistent with previous proteomic evidence of a delayed differentiation program. In contrast, MECP2^KO^ cells showed no deviation from WT at these demethylated sites on d4, indicating normal early demethylation. Sites that became hypermethylated from d0 to d4 in WT did not differ between WT and either KO. We then analyzed the top 10,000 CpG sites that were differentially methylated in WT cells during late differentiation (d4 to d8), including both hyper- and hypomethylated sites (Fig. 4C, Supplementary Table 3). These sites showed no genotype-dependent differences at d0 but began to diverge by d4. For sites that would later undergo demethylation in WT, all genotypes had similar methylation levels at d4. In contrast, sites destined to gain methylation in WT were hypermethylated in TET3^KO^ and hypomethylated in MECP2^KO^ at d4, therefore showing an inverse pattern between the two KOs. By d8, however, this divergence between the KOs resolved: sites demethylated in WT from d4 to d8 were more methylated in both KOs, while sites normally hypermethylated in WT were less methylated in both KOs (Fig. 4C, Supplementary Table 3).

Together, these data indicated that all genotypes start from a similar methylation state at d0, diverge with distinct patterns by d4, and the TET3^KO^ and MECP2^KO^ converge again at d8, where they exhibit highly similar methylation profiles that differ from WT. This further supports a shared trajectory between the KOs for the early neuronal maturation phase, despite their initially divergent methylation regulation mechanisms.

To investigate whether the convergence of methylation patterns in TET3^KO^ and MECP2^KO^ by d8 is also reflected at the level of gene expression, we next analyzed transcriptomic changes across the differentiation trajectory from d0 to d4 and to d8 in all three genotypes (Supplementary Table 4). PCA of the transcriptome revealed a pattern similar to the DNA methylation data, except that at d4 the genotypes did not segregate into distinct subclusters. By d8, however, WT samples were again clearly separated from both KOs, which in turn clustered closely together, indicating a converging transcriptomic profile in TET3^KO^ and MECP2^KO^ iNGNs (Fig. S5A). To assess how gene expression dynamics evolved during differentiation in TET3^KO^ and MECP2^KO^ iNGNs compared to WT, we first generated Alluvial plots tracking genes across two transitions: from d0 to d4, and from d4 to d8 (Fig. 4D, Supplementary Table 4). Genes were grouped according to whether their expression was significantly upregulated, downregulated, or unchanged in each interval. Overall, the expression trajectories, i.e., how genes transitioned between expression states from d0 to d4 and then d4 to d8, were highly similar across all three genotypes, indicating that the global structure of gene expression changes during iNGN differentiation is largely conserved. However, closer inspection revealed a distinct difference in one specific flow: a subset of genes that were downregulated from d0 to d4 and subsequently upregulated from d4 to d8 was more prominent in WT than in either KO. This gene set showing downregulation from d0 to d4 in all genotypes, but partial reactivation from d4 to d8 specifically in WT cells, is enriched for regulators of developmental signaling, lineage plasticity, and differentiation checkpoints (e.g., *POU5F1*, *DNMT3B*, *SOX21*, *OTX2*, *GLI3*, *CDH1*, *TRPC6*, *EPHA1*). For *POU5F1*, we already observed a hypermethylation status in both KOs compared to the WT (Supplementary Table 3), consistent with their transcriptional profile (Supplementary Table 4). Many of these factors normally act to restrain premature neuronal differentiation or maintain cellular adaptability. Their reactivation in WT at d8 likely reflects a transient plasticity window that facilitates neuronal maturation and circuit formation. In contrast, the continued downregulation in TET3^KO^ and MECP2^KO^ suggests a failure to re-engage this plasticity program, consistent with reduced transcriptional and cellular adaptability at later differentiation stages.

Next, we compared gene expression differences between the KOs and WT at each differentiation stage. At d0, all genotypes showed minimal transcriptional differences, consistent with low expression of TET3 and MECP2 in the pluripotent state (Fig. S5B, Supplementary Table 4). However, with ongoing differentiation, transcriptomic divergence increased (Fig. S5C, Supplementary Table 4). By d8, both KOs exhibited substantially altered gene expression profiles, with 2,551 differentially expressed genes (DEGs) in TET3^KO^ and 2,138 in MECP2^KO^ compared to WT, of which 1,716 genes were shared, representing a highly significant overlap (Fig. S5D). Moreover, fold changes relative to WT were strongly correlated between the two KOs, indicating that the transcriptional effects of *TET3* or *MECP2* deficiency converge also at the global transcriptome level by d8 (Fig. 4E, Fig. S5E, Supplementary Table 4). When we checked the hmC profile of selected genes at d8, we observed that the MECP2^KO^ featured a disrupted correlation between gene expression and hmC. For example, hmC levels in the gene body of *DNMT3B* were decreased in the TET3^KO^, but increased in the MECP2^KO^ compared to WT. However, transcriptional levels of DNMT3B were significantly reduced for both KOs (Fig. 4F). Functional enrichment analysis of DEGs commonly upregulated in both KOs revealed specific enrichment for cellular component GO terms associated with neuronal structural elements, including *neuron projection*, *synapse*, *axon*, *synaptic membrane*, and *dendrite* (Fig. 4G, Supplementary Table 4). Although a greater number of genes were commonly downregulated, and to a higher magnitude, than upregulated in the KOs, functional enrichment among downregulated DEGs was more diffuse. These genes were associated with broadly defined cellular component GO terms related to cell–cell communication (e.g., *cell junction*, *receptor complex*) and chromatin-associated structures (e.g., *chromosomal part*, *chromatin*) (Supplementary Table 4). Overall, our transcriptome data at d8 suggest highly similar impairments in transcriptional regulation and aberrations of neuronal function in both KOs despite inverse hmC patterns.

We next investigated whether the phenotypic convergence of TET3^KO^ and MECP2^KO^ neurons at d8 extends to the proteome (Supplementary Table 2). PCA revealed tight clustering of biological replicates within each genotype at both d0 and d4 (Fig. S5F). At these time points, WT and TET3^KO^ samples were clearly separated from MECP2^KO^ samples. Unlike the DNA methylation and transcriptome data, where PC1 primarily separated pluripotent from differentiated states, PC1 in the proteome data captured temporal progression from d0 to d4 to d8. By d8, all samples were closer in PCA space, but TET3^KO^ and MECP2^KO^ were indistinguishable from one another and remained distinct from WT (Fig. S5F). Comparison of differentially expressed proteins in the KOs relative to WT revealed an exceptionally high correlation between TET3^KO^ and MECP2^KO^ at d8 (Fig. 4H, Fig. S5G, Supplementary Table 2), further confirming their convergence at the proteomic level. As observed in the transcriptome, commonly downregulated proteins did not show specific GO term enrichment. In contrast, commonly upregulated proteins were enriched for transcriptional repressors, consistent with the predominance of downregulated transcripts in both KOs. Additionally, upregulated proteins showed strong enrichment for components involved in neuronal structure and connectivity, including multiple semaphorins, which are key regulators of axonal growth, neuronal polarity, neuronal migration, and synapse formation^59^, resulting in a GO term enrichment of processes involved in negative regulation of nervous system development, e.g. neuron differentiation or axon extension and guidance (Fig. S5H, Supplementary Table 2). On the other hand, enrichment was observed for proteins positively regulating synaptic transmission (Fig. S5H, Supplementary Table 2), such as synuclein-α and neuroligins. These proteomic changes translated into structural phenotypes as indicated by image-based quantification of synaptic vesicle density, which revealed a significant increase in both TET3^KO^ and MECP2^KO^ compared to WT, with no detectable difference between the two KOs (Fig. 4I, Fig. S5I).

In conclusion, although TET3^KO^ and MECP2^KO^ display inverse global hmC signatures, their cellular phenotypes and molecular profiles converge progressively during differentiation, resulting in remarkably similar DNA methylation, transcriptomic, and proteomic landscapes by d8.

## Discussion

This study identifies TET3 as a key regulator of neuronal maturation. Using iNGN-differentiation as a model, we show that *TET3* deficiency delays the progression of post-mitotic neurons toward a mature state, as evident from epigenetic, transcriptomic, proteomic, and functional analyses. During neuronal maturation, TET3 physically interacts with the methylation reader MECP2, whose loss causes Rett syndrome^24–26^. Although MECP2 antagonizes TET3 activity and both proteins display inverse hmC dynamics early during differentiation, their respective knockouts ultimately converge on similar transcriptional and structural phenotypes at late stages, suggesting a shared defect in chromatin adaptability.

Mechanistically, our data indicate that TET3 and MECP2 act at opposing yet interconnected levels of DNA modification–dependent chromatin regulation. mC in promoter and regulatory regions promotes chromatin compaction and transcriptional repression, while hmC supports chromatin accessibility and enhancer activity^60–62^. MECP2, binding both mC and hmC with different affinities, stabilizes nucleosome organization and higher-order chromatin architecture^53, 63, 64^. Our data suggest that the balance between TET3-mediated hydroxymethylation and MECP2-mediated chromatin modification ensures that neuronal chromatin remains both structurally stable and transcriptionally responsive, a prerequisite for differentiation and plasticity.

An additional layer of regulation may involve DNMT3A, the major de novo methyltransferase in post-mitotic neurons and another interactor of MECP2 with consequences for chromatin dynamics^37–40, 65^. Its stable expression in WT but premature loss in both KOs suggests an interdependent regulatory axis among DNMT3A, TET3, and MECP2, coordinating methylation turnover and chromatin organization during neuronal maturation. Disruption of this axis likely compromises the transient reactivation of developmental gene programs, resulting in reduced transcriptional flexibility at late differentiation stages.

Overall, our findings suggest that the dynamic balance between DNA methylation and hydroxymethylation defines cellular plasticity during differentiation and that disruption of this interplay may represent a shared pathogenic mechanism in neurodevelopmental disorders. Future work should address how TET3–MECP2 interactions are temporally regulated and how they intersect with DNMT3A-mediated methylation to shape higher-order chromatin structure in post-mitotic neurons. Such studies will be critical to understand how epigenetic plasticity supports neuronal maturation and function.

## Acknowledgements

FR Traube, T Carell, J Walter and S Michalakis thank the Deutsche Forschungsgemeinschaft (DFG) for financial support via CRC1309 (Project Nr. 325871075, Projects A04 (T Carell), A05 (J Walter), B05 (S Michalakis) and C08 (FR Traube). FR Traube and S Michalakis thank the DFG for financial support via SPP2502 EPIADAPT (Project Nr. 563452213). S Michalakis further acknowledges the support by the Münchener Universitätsgesellschaft e.V.. V Busskamp acknowledges support from the Volkswagen Foundation (Freigeist-A110720), the German Federal Ministry for Economic Affairs and Climate Action through the German Space Agency at DLR in the framework of SANSRETINA (50WB2516), the Deutsche Forschungsgemeinschaft (DFG) – Project Nr. 531985111 and the Cluster of Excellence - ImmunoSensation2 EXC-2151-390873048.

We thank Dr. Pavel Kielkowski and his team at the LMU Institute of Chemical Epigenetics for Eclipse maintenance, calibration and troubleshooting. We thank the Biomedical Center Munich Core Facility Flow Cytometry (LMU) for their support. We thank Kerstin Kurz (LMU), Kerstin Skokann (LMU) and Hafsa-Afandi Lee (Universität Stuttgart) for technical assistance.

## Author Contributions

All authors have reviewed, edited and approved the manuscript.

FR Traube: Conceptualization, Data curation, Formal analysis, Funding acquisition, Investigation (cell culture, MS experiments, coIPs, HEK293T assays, immunoblotting), Methodology, Project administration, Supervision, Visualization and Writing – Original draft.

G Gasparoni: Data curation, Formal analysis, Investigation (EPIC array and RNAseq), Methodology, Validation and Visualization.

A Winkler: Formal analysis, Investigation (cell culture, imaging experiments, qRT-PCRs), Methodology, Validation and Visualization.

AS Geserich: Investigation (cell culture, generation of MECP2^KO^ iNGNs).

H Sepulveda: Formal analysis, Investigation (CMS-IP), Methodology and Visualization.

JC Angel: Formal analysis and Methodology (CMS-IP).

X Yue: Investigation (CMS-IP).

R Habibey: Formal analysis, Investigation (MEA electrophysiology), Methodology and Visualization.

V Splith: Investigation (cell culture, generation of TET3^KO^ iNGNs, qRT-PCRs).

GI Gökҫe: Formal analysis, Investigation (cell culture, imaging experiments), Methodology, Validation and Visualization.

G Giorgio: Formal analysis, Investigation (cell culture, imaging experiments), Methodology and Visualization.

C Bernardini: Investigation (cell culture, MS experiments).

R Sachsse: Investigation (imaging experiments).

C Scheel: Investigation (cell culture, qRT-PCR).

M Bickerstaff-Westbrook: Methodology

M Biel: Funding acquisition and Resources.

T Carell: Funding acquisition, Methodology and Resources.

V Busskamp: Funding acquisition, Methodology, and Resources.

A Rao: Funding acquisition, Methodology and Resources.

J Walter: Conceptualization, Formal analysis, Funding acquisition, Methodology, Project administration, Resources and Supervision.

S Michalakis: Conceptualization, Formal analysis, Funding acquisition, Methodology, Project administration, Resources and Supervision.

## Data Availability

All datasets are available from the corresponding authors upon reasonable request. All omics datasets generated and analyzed in this study (EPIC-Array, RNAseq, CMS-IP and proteomics data) will be made publicly available in a dedicated, publicly accessible repository (e.g. NIH GEO^66^ for next-generation sequencing data and PRIDE^67^ for proteomics data) upon publication in a peer-reviewed journal. Other data types will be made available via general-purpose repositories such as *Zenodo* or *Figshare*.

## Competing Interests

The authors declare that they do not have any competing interests.

## Materials and Methods

### Cell Culture Procedures

All mammalian cell lines were maintained at 37 °C in a humidified atmosphere containing 5% CO₂. All cell handling was performed under sterile conditions. Routine testing for Mycoplasma contamination was carried out using a PCR-based detection kit (Jena Bioscience PP-401).

#### iNGNs

iNGN cells were routinely maintained in StemFlex™ Medium (Gibco™ A3349401) supplemented with 1% (v/v) Antibiotic-Antimycotic (Gibco™ 15240062) (referred to as stem cell medium).^46^ Prior to seeding, standard tissue culture plates were coated overnight (37 °C, 5% CO_2_, humidified atmosphere) with Geltrex™ LDEV-free, hESC-qualified basement membrane matrix (Gibco™ A1413302), diluted 1:50 in DMEM/F-12 (1:1) containing L-glutamine and 15 mM HEPES (Gibco™ 11330057), and supplemented with 1% Antibiotic-Antimycotic.

Cells were passaged upon reaching 70 – 80% confluency. For passaging, iNGNs were washed with Dulbecco’s phosphate-buffered saline (DPBS; Gibco™ 14190094), detached using TrypLE™ Express (Gibco™ 12604013), and collected in DPBS. After centrifugation (5 min, 300 × g), the cell pellet was resuspended as a single-cell suspension in StemFlex medium supplemented with 1% (v/v) RevitaCell™ Supplement (Gibco™ A2644501) and replated in a 1:5 ratio. Medium was replaced after 18 – 24 h with fresh StemFlex medium and changed every other day thereafter.

Neuronal induction was initiated by addition of doxycycline hyclate (Sigma-Aldrich D9891) at a final concentration of 0.5 µg/mL. This time point was designated as day 0 (d0). From d0 to d2, the culture medium was gradually transitioned from stem cell medium to neuronal medium consisting of Neurobasal™-A Medium (Gibco™ 10888022) supplemented with 1% (v/v) Antibiotic-Antimycotic, 1% (v/v) GlutaMAX™ (Gibco™ 35050038), 2% (v/v) B-27™ Supplement (Gibco™ 17504044), and 0.5 µg/mL doxycycline. On d1, the medium was replaced with a 1:1 mixture of stem cell and neuronal medium. From d2 onward, cells were maintained in neuronal medium with medium changes every other day until d8.

#### HEK293T

HEK293T cells (ATCC CRL-3216) were cultured in RPMI-1640 medium (Sigma-Aldrich R0883) supplemented with 10% (v/v) fetal bovine serum (Invitrogen 10500-064), 1% (v/v) L-alanyl-L-glutamine (Sigma-Aldrich G8541), and 1% (v/v) penicillin–streptomycin (Sigma-Aldrich P0781) (hereafter referred to as HEK293T medium). Cells were passaged routinely upon reaching 70 – 80% confluency.

For passaging, cells were washed with DPBS (Gibco™ 14190094), detached using TrypLE™ Express (Gibco™ 12604013), and collected in HEK293T medium. After centrifugation (5 min, 300 × g), the cell pellet was resuspended as a single-cell suspension in fresh medium and replated at a 1:10 ratio. Culture medium was replaced every other day.

### Generation of TET3^KO^ and MECP2KO iNGNs by CRISPR-Cas9 Genome Editing

*TET3* and *MECP2* knockout (KO) iNGNs were generated using CRISPR-Cas9-mediated genome editing. Guide RNAs (gRNAs) were designed to target early exons in the human *TET3* and *MECP2* genes, resulting in deletions of exon 7 and 8 in *TET3* and exon 3 and 4 in *MECP2*, respectively. The deletion in *TET3* introduced a frameshift and premature stop codon, disrupting the catalytic domain. The deletion in *MECP2* removed the methyl-CpG binding domain (MBD) and transcriptional repression domain (TRD), abolishing protein function.

Genomic and transcript sequences were obtained from the UCSC Genome Browser (GRCh38/hg38), NCBI Gene database, and Ensembl Genome Browser (release 93). gRNA sequences were identified using the CRISPOR tool with SpCas9 PAM setting (NGG) and evaluated for specificity and off-target potential.^68, 69^ gRNAs with specificity scores <50 were excluded. The following high-specificity gRNAs were used:

TET3 gRNAs

~~~
gRNA1: 5′-CACATGGGTTTACGGAAGTA-3′
~~~

~~~
gRNA2: 5′-CAACCCAAGTGCCCCTAACT-3′
~~~

~~~
gRNA3: 5′-CAGGTAGCCTGGTGTAAAAA-3′
~~~

~~~
gRNA4: 5′-TATCGCTTTACTTCTGGACT-3′
~~~

MECP2 gRNAs

~~~
gRNA1: 5′-GACAAGAGGCCGTCGACTGC-3′
~~~

~~~
gRNA2: 5′-CCTTGAAGTGCGACTCATGC-3′
~~~

~~~
gRNA3: 5′-GCACTGATGGCACCGAAAAC-3′
~~~

~~~
gRNA4: 5′-CTCTGTCTCTAACGACCACA-3′
~~~

gRNAs were delivered as ribonucleoprotein (RNP) complexes using the Alt-R® CRISPR-Cas9 system (Integrated DNA Technologies). RNP complexes were assembled from Alt-R® crRNAs, tracrRNA, and Cas9 Nuclease V3 according to the manufacturer’s protocol. Complexes were introduced into iNGN cells via nucleofection using the Amaxa™ Human Stem Cell Nucleofector® Kit 2 (Lonza VPH-5022) and the Nucleofector™ 2b device (Lonza).

Following nucleofection, cells were cultured under standard iNGN conditions for recovery. Single-cell clones were isolated by fluorescence-activated cell sorting (FACS) at the Flow Cytometry Core Facility, Biomedical Center, LMU Munich. Dead cells were excluded using propidium iodide, and viable clones were expanded for genotyping.

To confirm gene disruption, genomic DNA (gDNA) was isolated from expanded clones, and PCR was performed using primers flanking the targeted regions:

TET3^KO^ genotyping primers

~~~
Forward: 5′-CTGCCCTTATCAGAGTTTTTCCCAT-3′
~~~

~~~
Reverse: 5′-GCCTCACCAGTGGATACCAG-3′
~~~

Amplified sequences: WT – 4601 bp; TET3^KO^ - 852 bp

MECP2^KO^ genotyping primers

~~~
Forward: 5′-TTCATGTTTGGAAAGCGGCA-3′
~~~

~~~
Reverse: 5′-CTCAAGGGACGTCCTCCAAC-3′
~~~

Amplified sequences: WT – 3509 bp; MECP2^KO^: 648 bp

PCR reactions were carried out using Q5® High-Fidelity DNA Polymerase (New England Biolabs M0491L). PCR products were visualized by agarose gel electrophoresis, and successful deletions were confirmed by Sanger sequencing (Eurofins Genomics).

### Imaging

#### Immunofluorescence Staining

For immunofluorescence, iNGNs were seeded on Geltrex™-coated glass coverslips and cultured under standard conditions. All staining steps involving fluorescent dyes were performed under light-protected conditions.

Prior to staining, coverslips were washed with DPBS and fixed with 4% formaldehyde (Sigma-Aldrich P6148) in DPBS for 20 min at room temperature, followed by additional DPBS washes. Cells were then permeabilized and blocked for 1 h in blocking buffer consisting of 2% (w/v) BSA (Gibco™ 30066575) and 0.5% (v/v) Triton X-100 (Sigma-Aldrich 648466) in DPBS.

Primary antibodies diluted in blocking buffer were applied and incubated overnight at room temperature in a light-protected humidified chamber. The next day, cells were washed twice with DPBS and incubated with fluorophore-conjugated secondary antibodies, also diluted in blocking buffer, for 2 h at room temperature. After two additional DPBS washes, cell nuclei were stained with 1:1000 diluted DAPI (Invitrogen^TM^ 62248) for 10 min. Coverslips were washed again, mounted onto glass slides using Aqua-Poly/Mount (Polysciences 18606), and dried overnight.

#### BrdU Incorporation Assay for Immunofluorescence

To label S-phase cells, BrdU (Sigma-Aldrich B5002) was added to the culture medium at a final concentration of 0.3 µg/mL for 3 h. Cells were then washed five times with DPBS and fixed as described above. For DNA denaturation, fixed cells were incubated with 2 N HCl for 45 minutes, followed by neutralization with 0.1 M borate buffer (pH 8.5) for 30 minutes. Subsequent staining steps were performed as outlined above.

#### Antibodies for Immunofluorescence

Primary antibodies

- Rat anti-BrdU (Abcam, ab6326; 1:200)
- Rabbit anti-Ki67 (Invitrogen, MA5-14520; 1:500)
- Mouse IgG1 anti-SV2A (DSHB, AB_2315387; 1:10)
- Mouse IgG2a anti-TUBB3 (TUJ1, BioLegend, 801213; 1:300)

Secondary antibodies

- Donkey anti-rat Cy3 (Jackson ImmunoResearch, 712-165-153; 1:500)
- Goat anti-rabbit Alexa Fluor 488 (Cell Signaling, 4412S; 1:500)
- Goat anti-mouse IgG1 Alexa Fluor 488 (Invitrogen, A21121; 1:1000)
- Goat anti-mouse IgG2a Alexa Fluor 647 (Invitrogen, A21241; 1:500)

#### Immunofluorescence and Brightfield Microscopy Image Acquisition

Representative brightfield images of iNGNs were acquired using the EVOS® FL Cell Imaging System (Life Technologies) under standardized acquisition settings across all samples.

Immunofluorescence imaging was performed on an inverted Leica SP8 confocal microscope (Leica Microsystems) equipped with lasers at 405, 488, 522, and 638 nm. Images were acquired using a HC PL APO 40x/1.30 Oil CS2 objective with type F immersion liquid (Leica Microsystems) and LAS X software version 3.5.1.18803 (Leica Microsystems).

Post-acquisition image processing and quantitative analyses were performed using Fiji (ImageJ) software^70^. For every marker, imaging settings including gain and laser power as well as image processing settings were kept constant among different samples and corresponding controls. Immunofluorescence-labeled cells were manually quantified using the multi-point tool of Fiji Image J Software. DAPI-stained nuclei served as a reference for total cell numbers.

#### Quantification of iNGN Cell Counts from Brightfield Images

Cell number quantification was performed using Cellpose 2.0 GUI^71^. To enable fast and reproducible cell counting across different morphological stages of neuronal differentiation, three custom Cellpose models^72^ were trained for iNGNs at d0, d4, and d8. Models were developed using 10× magnification images acquired with the EVOS® FL Cell Imaging System (Life Technologies), which were subdivided into four quadrants (equivalent to 20×) to facilitate improved training efficiency. Each model was trained on nine quadrant images (three per biological replicate) using default Cellpose parameters. Regions of interest (ROIs) were manually corrected during training to refine segmentation. Given the minimal phenotypic differences among genotypes at the morphological level, model training was conducted exclusively on WT iNGN images. The final models were then applied to all genotypes for their respective time points. Cell counts were obtained directly from the automated model output, without subsequent manual correction.

#### Quantification of Synaptic Vesicles per Neurite Length

Synaptic vesicles were quantified based on SV2A immunostaining. The SV2A channel was processed using Gaussian blur (σ = 1) followed by the “Find Maxima” function (prominence = 7) in Fiji. Detected maxima were overlaid onto the nuclear (DAPI) channel to identify soma-associated vesicles; only vesicles within clearly defined somata were included. To assess synaptic vesicle density along neurites, the total neurite length was quantified based on TUBB3 staining using a custom Fiji macro. The macro executed the following steps: (1) channel splitting, (2) contrast enhancement (saturated = 0.35), (3) auto-thresholding using the “Percentile dark” method, (4) conversion to binary mask, (5) skeletonization, and (6) analysis via “Analyze Skeleton (2D/3D)” with no pruning. The resulting “Skeleton Branch Info” output was used to calculate total neurite length per image. Synaptic vesicle density was then calculated as the number of soma-associated SV2A puncta per micrometer of neurite length. Quantification was performed across all biological and technical replicates.

### Isolation of Nucleic Acids and Proteins

#### Nucleic Acids

Total RNA from iNGNs for qRT-PCR was extracted using the RNeasy Plus Kit (Qiagen 74134) according to the manufacturer’s protocol.

For EPIC-Array, next-generation sequencing and MS approaches, gDNA and RNA were isolated as previously described^28^.

#### Protein Isolation

For proteomics, ca. 1 ×10^6^ cells were lysed in 200 µL of total lysis buffer (20 mM HEPES, 1% (v/v) NP-40, and 0.2% (w/v) SDS) for 30 min on ice and afterwards centrifuged at 21,000g at 4 °C for 10 min. The supernatant containing the proteins was transferred to a new tube. Total protein concentration was determined using a BCA assay (Thermo Scientific^TM^ Pierce^TM^ A55864) according to the manufacturer’s protocol.

For the preparation of nuclear extracts, ca. 4 × 10^6^ cells were harvested and nuclear extracts were prepared as previously described^73^ with the modification that every buffer was supplemented with cOmplete^TM^ protease inhibitor (Roche 04693132001) instead of PMSF to inhibit any protease activities. Concentration of nuclear proteins were determined using a BCA assay. Enrichment of nuclear proteins compared to total proteome was tested by LC-MS/MS as previously described^74^.

For total cell lysis with subsequent coIP, ca. 1 × 10^6^ cells were lysed in 250 µL gentle lysis buffer (20 mM Tris pH = 8.0, 137 mM NaCl, 2 mM EDTA, 5 mM EGTA, 1% (v/v) NP-40, 1×cOmplete^TM^ protease inhibitor + 500 U (2 µL) Benzonase nuclease (Millipore E1014-5KU)) and incubated on a rotating wheel for 1 h at 4 °C. Afterwards, the lysate was centrifuged for 15 min at 21,000 × g at 4 °C and afterwards, the supernatant was transferred to a new tube and protein concentration was determined using a BCA assay.

### Quantitative Real-Time PCR (qRT-PCR)

cDNA synthesis was performed using the RevertAid First Strand cDNA Synthesis Kit (Thermo Scientific K1622) following the manufacturer’s instructions. For qRT-PCR, cDNA was diluted 1:5 with nuclease-free water.

qRT-PCR was conducted using either a StepOnePlus Real-Time PCR System or a QuantStudio 5 Real-Time PCR System (Applied Biosystems), in combination with the PowerUp SYBR Green Master Mix (Applied Biosystems A25742), according to the manufacturer’s instructions. Each reaction was run in two technical replicates per gene per sample. Gene expression was normalized to the housekeeping gene ACTB, and relative expression levels were calculated using the ΔC_t_ method^75^ assuming a primer efficiency of 2.

For *TET1*, *TET2*, and *TET3*, expression values were further normalized to *TET1* expression at d0. Baseline settings and threshold values were manually adjusted using the corresponding system software (StepOnePlus or QuantStudio Design and Analysis Software v1.5.2, Applied Biosystems).

Gene-specific primers were designed using NCBI’s Primer-BLAST tool, with a GC content of 40 – 60% and self-complementarity scores below 4. The primer sequences used in this study are as follows:

~~~
ACTB forward: 5’-GCCGCCAGCTCACCAT-3’
~~~

~~~
ACTB reverse: 5’-CACGATGGAGGGGAAGACG-3’
~~~

~~~
ETT1 forward: 5’-GCTCTCATGGGTGTCCAATTGCT-3’
~~~

~~~
TET1 reverse: 5’-ATGAGCACCACCATCACAGCAG-3’
~~~

~~~
TET2 forward: 5’-AAGGCTGAGGGACGAGAACGA-3’
~~~

~~~
TET2 reverse: 5’-TGAGCCCATCTCCTGCTTCCA-3’
~~~

~~~
TET3 forward: 5’-GCAAGACACCTCGCAAGTTC-3’
~~~

~~~
TET3 reverse: 5’-CCTCGTTGGTCACCTGGTTC-3’
~~~

### Mass spectrometry

#### QQQ-MS

QQQ-MS for absolute quantification of DNA modifications, including downstream analysis, were performed as previously described^28^.

#### Proteomics (LFQ-DIA – Whole Proteome)

##### SP3 protocol

For each sample, 20 μg of protein was processed. The protein lysate was added to pre-washed Dynabeads™ Protein G (Invitrogen™ 10003D) at a bead-to-protein ratio of 10:1 (approximately 7 μL resuspended beads per 200 μg protein), and the volume was adjusted to 50 μL using total lysis buffer. Samples were incubated with the beads in an Eppendorf ThermoMixer C at room temperature with shaking at 1,000 rpm for 1 min. Following incubation, 120 μL of ethanol (EtOH; Fisher Chemicals A4564FB25) was added, and samples were incubated for an additional 5 min at 1,000 rpm. Beads were immobilized using a magnetic rack, and the supernatant was discarded. Bead-bound proteins were washed three times with 100 μL of 80% (v/v) EtOH, each time shaking at 850 rpm for 1 min, followed by magnetic separation and removal of the supernatant. Beads were resuspended in 100 μL of 100 mM ammonium bicarbonate (ABC; Sigma-Aldrich A6141). Protein reduction and alkylation were performed by sequential addition of 10 mM DL-dithiothreitol (DTT; Sigma-Aldrich 10708984001) and 20 mM iodoacetamide (IAA; Sigma-Aldrich I1149) from 1 M stock solutions. Samples were incubated at 95 °C for 5 min with shaking at 850 rpm. After cooling to room temperature, trypsin (LC-MS grade; Thermo Scientific™ Pierce™ A40009) was added at a 1:50 enzyme-to-protein ratio (0.4 μg trypsin per 20 μg protein), and digestion was carried out overnight at 37 °C with shaking at 850 rpm. The resulting peptides were transferred to a fresh tube, and the beads were washed twice with 50 μL of 0.1% (v/v) formic acid (FA; Optima™ Fisher Chemical A11750). All fractions were pooled and cleared of residual beads by magnetic separation. Peptide samples were stored at −80 °C until LC-MS/MS analysis.

##### MS acquisition and analysis

Mass spectrometry was performed using an Orbitrap Eclipse Tribrid mass spectrometer (Thermo Fisher Scientific™) equipped with a FAIMS interface and coupled to an UltiMate 3000 Nano HPLC system (Thermo Fisher Scientific™) via an EASY-Spray source (Thermo Fisher Scientific™). Data were acquired in data-independent acquisition (DIA) mode, and all solvents used were LC-MS grade. For each sample, 300 ng of peptides was first loaded onto an Acclaim PepMap 100 μ-precolumn cartridge (5 μm, 100 Å, 300 μm ID × 5 mm; Thermo Fisher Scientific™), followed by separation on an in-house–packed analytical column consisting of a PicoTip emitter (non-coated, 15 cm length, 75 μm ID, 8 μm tip; New Objective) packed with Reprosil-Pur 120 C18-AQ material (1.9 μm, 150 Å; Dr. A. Maisch GmbH). The column was maintained at 40 °C. Peptides were separated using a 60-minute linear gradient at a flow rate of 0.3 μL/min with buffer A (0.1% formic acid in water) and buffer B (0.1% formic acid in acetonitrile) as follows:

- 0 – 5 min: 4% B
- 5 – 6 min: ramp to 7% B
- 6 – 36 min: linear increase to 24.8% B
- 36 – 41 min: ramp to 35.2% B
- 41 – 41.1 min: rapid increase to 80% B
- 41.1 – 46 min: hold at 80% B
- 46 – 55 min: re-equilibration at 4% B

The DIA duty cycle included one MS1 scan followed by 30 MS2 scans. MS2 isolation windows were 4 *m/z* wide with 2 *m/z* overlaps. MS1 spectra were acquired in the Orbitrap at 60,000 resolution over a mass range of 200 – 1,800 *m/z,* with the RF lens set to 30%. MS2 spectra were acquired in the Orbitrap at 30,000 resolution, also with the RF lens at 30%, covering a precursor mass window of 500–740 m/z. Higher-energy collisional dissociation (HCD) was used for fragmentation with a fixed collision energy of 35%. FAIMS was enabled during the entire duty cycle, with a single compensation voltage (CV) of −45 V applied to both MS1 and MS2 scans.

##### Analysis of MS spectra and pathway analysis

Raw DIA-MS files were processed using DIA-NN (v. 1.8.1)^76^ to identify and quantify proteins across samples. The analysis was conducted in library-free mode with FASTA digest enabled, using a UniProt-derived *Homo sapiens* reference proteome. The following parameters were applied:

- Protease: Trypsin/P
- Allowed missed cleavages: 2
- Minimum peptide length: 6 amino acids
- Match Between Runs (MBR): Enabled
- Heuristic protein inference: Disabled
- Quantification strategy: Robust LC (high precision)
- Number of processing threads: 7

All other parameters were left at default settings.

The resulting protein group output was imported into Perseus (v2.0.11)^77^ for downstream analysis. Samples were grouped by genotype and differentiation day. To ensure robust quantification, only proteins detected in at least 75% of samples (≥3 out of 4 replicates) within at least one group were retained. Protein intensities were log_2_-transformed, and missing values were imputed from a normal distribution (column-wise, default settings in Perseus). Pairwise group comparisons were performed using the volcano plot function in Perseus (two-sided t-test, 250 randomizations, FDR = 0.05). Functional enrichment analyses for significantly differentially expressed proteins were conducted using Reactome^48^ and GOrilla^48^, with full details available in Supplementary Table *2*.

#### Proteomics (LFQ-DDA – coIP)

##### coIP of GFP-enriched TET3

The coIPs using recombinantly expressed GFP-TET3 for subsequent LC-MS/MS analysis were performed using iNGN d8 nuclear lysate and analyzed as previously described^73^. Recorded spectra were further analyzed using MaxQuant^78^ and Perseus^77^ with significance thresholds for enriched proteins set to −log(p-value) > 1.3 and difference (log_2_FC) > 1.

### Expression Plasmids

MECP2E1-MycDDK (plasmid with kanamycin resistance, CMV promoter, OriGene MR226839)

(N-terminus) MECP2-MycDDK; MECP2 (UniprotKB – Q9Z2D6-2) starting from amino acid 1 of the original sequence. MECP2E1 T158M threonine to methionine mutation at position 175; MECP2E1 R306C arginine to cysteine mutation at position 323.

GFP-TET3^49^ (plasmid with ampicillin resistance, CAG promoter):

(N-terminus) eGFP – TEV cleavage site – Tet3; murine Tet3 (UniprotKB – Q8BG87-4) starting from amino acid 1 of the original sequence.

#### Site-directed mutagenesis

To introduce the T148M or R306C mutations in the MECP2-MycDDK expression plasmid, PCR site-directed mutagenesis was used.

Primer for site-directed mutagenesis:

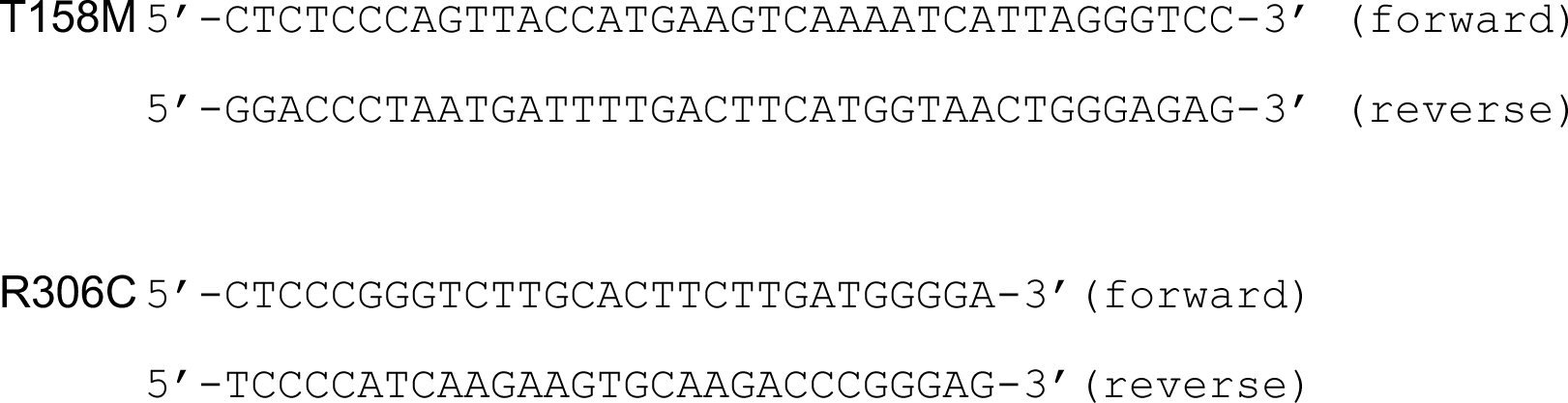

PCR reaction set up per sample:

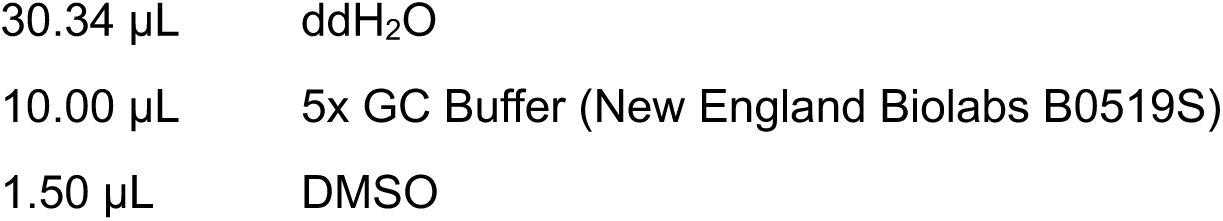
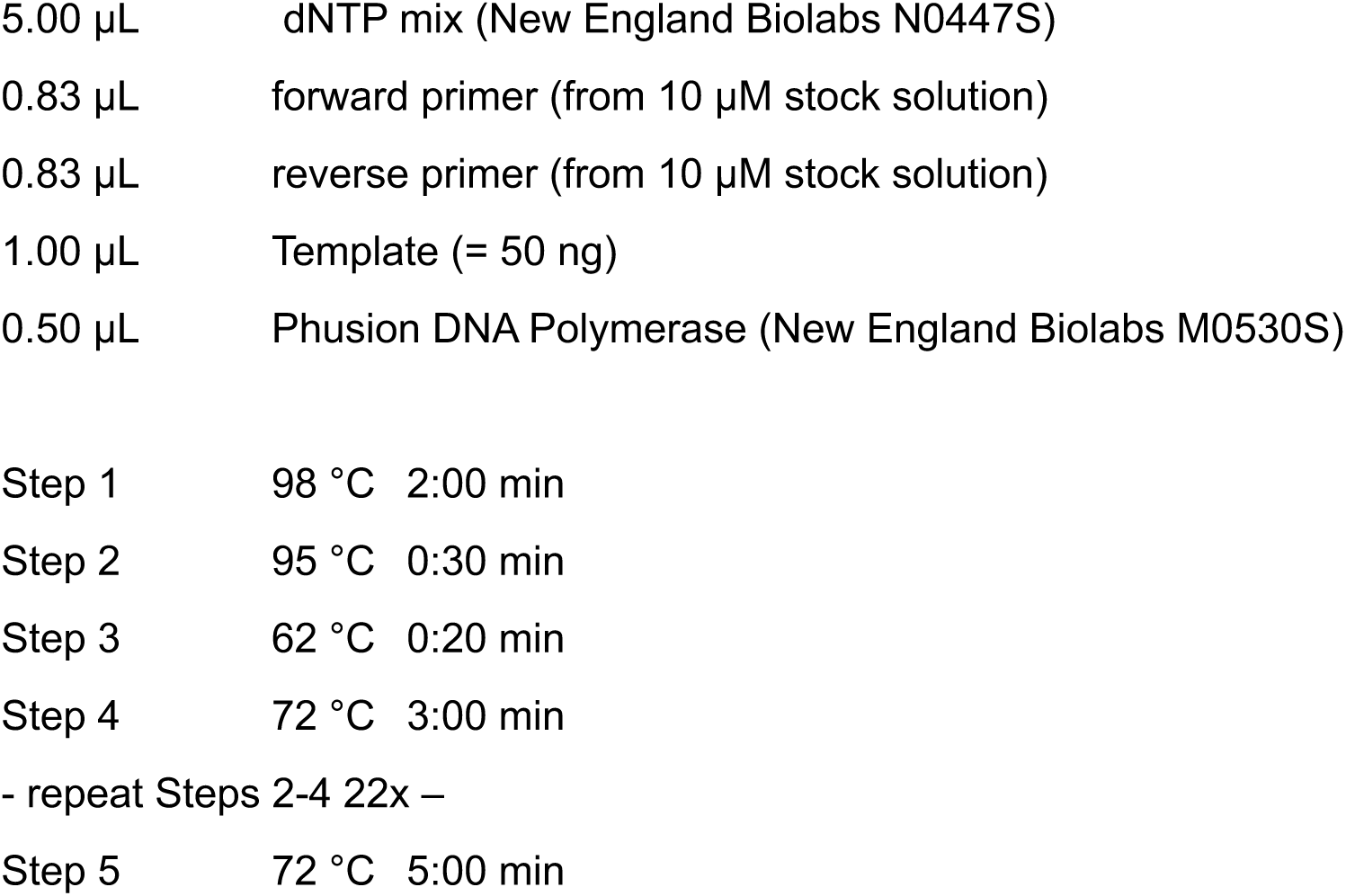

After the PCR, 10 µL of the PCR product was checked on an 1% agarose gel. The rest was stored on ice while the agarose gel was running. Agarose gel electrophoresis confirmed that in any case only one PCR product at the correct bp size was obtained. Subsequently, 1 µL of DpnI (New England Biolabs R0176S) was added per sample and samples were incubated for 1 h at 37 °C. Afterwards, the mix was used completely for transformation of chemical competent Escherichia coli (E. coli, New England Biolabs C2987I). Plasmids were isolated from positive clones and checked by Sanger sequencing for having the correct mutation.

### HEK293T Activity Assay

#### Transfection

2.5×10^5^ cells were seeded per well in 3 mL of RPMI medium. 24 h after seeding, the cells were transfected using 1.5 µg of DNA per expression plasmid (combination of GFP-TET3 and one of the MECP2-MycDDK expression plasmids), 4 µL of jetPRIME (Sartorius 101000015) and 150 µL of jetPRIME buffer. In case of GFP-TET3 only expressing cells, 1.5 µg of pESG-iba45 (iba 5-4445-001) were co-transfected to keep the amount of transfected DNA constant. Six hours after transfection, the medium was changed and 24 h after transfection, the cells were harvested and washed once with 1 mL of ice-cold DPBS.

#### Activity assay

100 µL of cell suspension were subsequently used for GFP signal quantification of living cells using flow cytometry on a BD LSRFortessa with the following settings: FSC 130 V, SSC 300 V, GFP 370 V log, GFP-A > 10^3^, 10000 events per measurement, the rest was lysed and gDNA was isolated and analyzed by UHPLC-QQQ-MS. The hmC/N values were subsequently divided by the GFP signal to obtain hmC/(N*RFU).

### CoIPs analyzed by Immunoblotting (Western blotting)

#### CoIP of endogenous TET3

CoIP of endogenous TET3 was performed using nuclear lysates from iNGNs at day 8 of differentiation. A total of 500 µg nuclear protein was incubated with 2 µg of anti-TET3 antibody (Abiocode R1092-N1, rabbit polyclonal) for 2.5 h at 4 °C on a rotating wheel. Subsequently, 20 µL of Dynabeads™ Protein G (Thermo Fisher Scientific, 10003D), pre-washed three times in 500 µL bead wash buffer (10 mM Tris-HCl pH=7.5, 150 mM NaCl, 0.5 mM EDTA), were added to each reaction and incubated for an additional 1 h at 4 °C under rotation. As a negative control, 500 µg of the same nuclear lysate was incubated with 2 µg of control goat IgG (Sigma-Aldrich G5518-2ML) and processed identically. After incubation, beads containing antibody-bound TET3 and its associated proteins were separated using a magnetic rack and washed three times with 500 µL CoIP wash buffer (10 mM HEPES pH=7.5, 150 mM NaCl, 0.5 mM EDTA) to remove non-specifically or weakly bound proteins.

Bound proteins were eluted by resuspension of the beads in 50 µL of SDS loading buffer (50 mM Tris-HCl pH=6.8, 100 mM DTT, 2% (w/v) SDS, 10% (v/v) glycerol, 0.1% (w/v) bromophenol blue) and incubation at 70 °C for 10 min. Eluates were subsequently analyzed by immunoblotting.

#### CoIP of recombinantly expressed GFP-TET3 and MECP2-MycDDK

For coIP of tagged proteins, 250 µg of total lysate from HEK293T cells transfected with GFP-TET3 and MECP2-MycDDK constructs or an empty vector control (as described in the HEK293T activity assay section) was used. Cells were lysed in gentle lysis buffer, and the lysates were incubated with 2 µg of anti-Myc antibody (Cell Signaling Technology, 2276S; mouse monoclonal, clone 9B11) to precipitate MECP2-Myc and associated proteins.

CoIP was performed following the same protocol as described for endogenous TET3 CoIP in iNGNs, including antibody incubation, bead capture using Dynabeads™ Protein G, washing steps, and elution in SDS loading buffer for subsequent analysis by immunoblotting.

#### SDS-PAGE and Western blotting

For SDS polyacrylamide gel electrophoresis (PAGE) with subsequent western blotting, protein samples in SDS loading buffer were resolved on a 4 - 15% precast polyacrylamide gel (4–15% Mini-PROTEAN® TGX™ Precast Protein Gels Biorad 4561083) using color-coded prestained protein standards (New England Biolabs P7712 or P7719) as molecular weight markers. For input samples from iNGN co-immunoprecipitation (coIP), 35 µg of total protein (corresponding to ∼7% of the total input used for coIP) were loaded. For input samples from coIP of transfected HEK293T cells, 10 µg (corresponding to ∼4% of input) were used. From eluted coIP fractions, 15 µL were loaded per lane and from total protein lysate to confirm absence of MECP2 in the MECP2^KO^ compared to WT 35 µg were loaded. Gels were run in SDS running buffer (25 mM Tris, 192 mM glycine, 0.1% (w/v) SDS) at a constant voltage of 150 V for 60 min.

For blotting, PVDF membranes (Amersham Hybond-P 0.45 µm, GE Healthcare, 10600023) were activated in methanol for 1 min, rinsed in water, and equilibrated in Towbin blotting buffer (25 mM Tris, 192 mM glycine, 20% (v/v) methanol, 0.038% (w/v) SDS) for 1 minute. Gels were also equilibrated in Towbin buffer for 5 min, and Whatman blotting papers (Sigma-Aldrich, WHA10426981) were equilibrated for 15 min. Wet/tank transfer was performed at 4 °C for 1.5 h at constant 45 V and for additional 3.5 h at constant 50 V.

Following transfer, membranes were blocked in 5% (w/v) non-fat milk in TBS-T (20 mM Tris pH=7.5, 150 mM NaCl, 0.1% (v/v) Tween-20) for 1 h at room temperature. Membranes were then incubated with primary antibodies diluted in 5 mL of 5% milk in TBS-T for 12 hours at 4 °C with gentle shaking. After incubation, membranes were washed 3 × 5 minutes with TBS-T, followed by incubation with HRP-conjugated secondary antibodies, diluted in 5% milk in TBS-T, for 1 hour at room temperature under gentle agitation. Membranes were then washed twice with TBS-T and once with TBS (TBS-T without Tween-20) and developed using SuperSignal™ West Pico Chemiluminescent Substrate (Thermo Scientific, 34077).

##### Primary and secondary antibodies for detection

Primary antibodies:

- Rabbit anti-GFP (Torrey Pines Biolabs TP401, Lot 081211, 1:1000)
- Rabbit anti-Histone H3 (Cell Signalling 4499S, clone D1H2, 1:1000)
- Rabbit anti-MECP2 (Diagenode C15410052, Lot A20-0042, 1:1000)
- Mouse anti-Myc tag (Cell Signalling 2276S, clone 9B11, 1:500)
- Rabbit anti-TET3 (Abiocode R1092-N1, Lot 7083, 1:250)

Secondary antibodies:

- HRP-conjugated anti-mouse IgG (Sigma-Aldrich AP130P, 1:5000)
- HRP-conjugated anti-rabbit IgG (Sigma-Aldrich A0545, 1:5000)

### EPIC Array and Next-Generation Sequencing Approaches

#### DNA Methylation Profiling via EPIC Array

For each sample, 500 ng of genomic DNA was subjected to bisulfite conversion using the EZ DNA Methylation Kit (Zymo Research D5002) according to the manufacturer’s instructions, with a final elution volume of 10 µL. Subsequently, 4 µL of bisulfite-converted DNA per sample was used as input for the Infinium MethylationEPIC BeadChip assay v1 (Illumina, San Diego), following the vendor’s protocol. Arrays were scanned on the Illumina HiScan platform.

Raw data preprocessing was performed using the RnBeads R/Bioconductor package (v2.12.2)^79,80^. Low-quality samples and probes were filtered using the *Greedycut* algorithm, based on a detection p-value threshold of 0.05. In addition, 17,371 CpG sites were excluded due to probe overlap with three or more annotated SNPs (*filtering.snp = 3*), and 2,983 non-CpG methylation sites were removed (*filtering.context.removal = CC, CAG, CAH, CTG, CTH, Other*). A further 111 CpG sites were filtered out for missing values in >80% of samples (*filtering.missing.value.quantile > 0.8*). In total, 20,465 sites were removed, resulting in a final dataset comprising 844,210 CpG sites across all samples.

Methylation levels at each CpG site were quantified as β-values, representing the proportion of methylated cytosines (ranging from 0 = unmethylated to 1 = fully methylated). β-values were normalized using the ‘dasen’ method from the *watermelon* package, as implemented in RnBeads.^81, 82^

Differential methylation analyses were performed via two-sided Student’s t-tests. Statistical signals were ranked using the *combined rank* method within RnBeads, which integrates the nominal p-value and the absolute mean methylation difference (delta-β) between groups. All downstream analyses and data visualizations were conducted in R.

#### mRNA Sequencing

For each sample, 1,000 ng of total RNA was used to generate sequencing libraries using the NEBNext® Ultra™ II Directional RNA Library Prep Kit with Beads (New England Biolabs E7765L) in combination with the NEBNext® Poly(A) mRNA Magnetic Isolation Module (New England Biolabs E7490L). Library preparation was performed according to the manufacturer’s protocol, including 8 PCR cycles for final amplification. Clean-up of the amplified libraries was carried out with 0.8× volume of AMPure XP beads (Beckman Coulter A63881), and libraries were eluted in 20 µL of EB buffer (Qiagen 19086).

Library concentration was quantified using the NEBNext Library Quant Kit for Illumina (E7630L), and fragment size distribution was assessed on an Agilent 2100 Bioanalyzer using the High Sensitivity DNA Kit (Agilent 5067-4626). Libraries were sequenced on an Illumina NextSeq 500 platform, yielding 25–50 million 75 nt single-end reads per sample.

Following demultiplexing and FASTQ file generation using the Illumina *bcl2fastq* tool, reads were processed using the grape-nf RNA-seq pipeline^83^ implemented in Nextflow v20.10.0. The pipeline incorporated the following tools: samtools^84^ version 1.3.1, RSEM^85^ version 1.2.21, STAR aligner^86^ version 2.4.0j, sambamba^87^ version 0.7.1, bamstats^88^ version 0.3.4 and RSeQC^89^ version 2.6.4. Reads were aligned to the human genome reference *GCA_000001405.15_GRCh38_no_alt_analysis_set* (NCBI) and annotated using GENCODE v22^90^. Gene-level quantification and differential expression analysis were performed in R using the DESeq2 package^91^. All downstream analyses and data visualization were conducted in R using the packages ggplot2 and ComplexHeatmap^92^.

#### CMS-IP

CMS-IP was performed as previously described^58^.

### MEA Electrophysiology and Optogenetic Stimulation

#### Cell culture

WT and TET3^KO^ iNGNs were passaged two times and cultured in Matrigel-coated well plates in mTeSR™1 medium (mTeSR™1 Basal Medium plus mTeSR™1 5x Supplement (STEMCELL^TM^ 85850) and 1% (v/v) penicillin-streptomycin). Doxycycline (0.5 μg/mL) was applied daily for five days for neural induction. At d1 after induction, we added 10 ul of the Lentiviral construct to initiate the expression of the EYFP tagged ChRimosn channels in all cell types. At d4 post-induction Ara-C (5 μM) was added to remove undifferentiated cells. Starting from d5 on, cells were re-seeded on MEA chips and media changed to BrainPhys™ Neuronal Medium (BrainPhys™ Neuronal Medium (STEMCELL^TM^ 05790) supplemented with 1% (v/v) penicillin-streptomycin, NeuroCult™ SM1 Neuronal Supplement (STEMCELL^TM^ 05790), N2 Supplement-A (STEMCELL^TM^ 05793), 20 ng/ml recombinant Human BDNF (STEMCELL^TM^ 78005), 20 ng/ml recombinant Human GDNF (STEMCELL^TM^ 78058) and 200 nM ascorbic acid. A day before re-seeding standard MEA chips were plasma treated and coated with Poly-D-lysine (PDL, 1 mg/ml stock, 50 µL per electrode area) and incubated overnight at 37°C. Extra PDL was washed out 3 times by sterile ddH2O and dried. Laminin was mixed in BrainPhys™ Media (0.05 mg/ml), added to the electrode area (50 µL) and incubated for three hours before re-seeding. All cell types were dissociated by Accutase, centrifuged (400 g, 4 min), re-suspended in Brainphys media and then were seeded on each MEA chip (100 K cells per MEA in total). To support the survival and electrophysiological properties of all neuronal cell lines we prepared them as banker culture system as following: rat primary cortical astrocytes (Thermo Fisher Scientific^TM^ A1261301) were cultured on coverslips with paraffin feets, worked as spacer, in astrocyte medium (DMEM plus N2 Supplement, 10 % (v/v) One Shot™ Fetal Bovine Serum and 1% penicillin-streptomycin) 5 days before re-seeding. After re-seeding the neural cells, astrocyte culture on coverslip was flipped on top of the neural cells on MEA substrate to provide banker culture system with continuous exposure of neurons to the astrocyte secreted materials. Every seven days half of the media was exchanged by fresh BrainPhys™ Media. Brightfield and fluorescent Images were acquired using the EVOS™ FL Imaging System equipped with RFP filter cube to detect neural cells expressing the EYFP tagged ChRimson channels.

#### Electrophysiology and optogenetic stimulation

Spontaneous neural network activity was measured by seeding the cells on standard MEA chips (Multi Channel Systems, 60 MEA200/30iR-Ti-gr) and acquisition of data using MEA1060-Inv-BC (sampling rate 25K Hz) and software user interface (MC_Rack) provided by Micro Channel Systems (MCS). Spontaneous activity of the differentiated neural networks were measured at days 15 and 25 post-induction (dpi). Optical stimulations with different pulse frequencies were applied at both recording days. Full-field optical stimuli (588 nm) at around 1 mW/mm2 intensity were applied using a Spectra4 lumencor LED light source through a 4x objective lens of an inverted Nikon microscope. Every optical stimulation experiment was composed of 3 phases including baseline recording of spontaneous activity for 3 minutes followed by yellow light pulses (588 nm) with 100 ms pulse width at different frequencies (0.2Hz, 0.5Hz, 1Hz and 5Hz) and different pulse numbers (9, 22, 49, and >49 pulses, respectively). Timestamps of the light stimulation pulses were recorded as digital data for later analysis.

#### Analysis of the electrophysiology data

Recorded data was replayed, and filtered (Butterworth 2nd order, high pass filter cut-off at 100 Hz) and timestamps of the spikes were detected by a negative or positive thresholds (−5 StDev of the peak-to-peak noise or +5 StDev of the peak-to-peak noise). Electrodes which showed response to light stimuli were considered as active electrodes and included in the statistical analysis. Action potential (AP) frequency was measured in light stimulated and non-stimulated periods at each individual electrode. For each experiment the average AP frequency was measured in baseline and during light stimulation periods. Timestamps of APs in each individual electrode was extracted in Neuroexplorer software.

### Statistical Analysis

If not described otherwise, statistical analysis was performed using GraphPad Prism v10.3.1. Details are given in Supplementary Table 1.

## Notes

### Competing Interest Statement

The authors have declared no competing interest.

