## Supplementary Figures for "Crosstalk between the Methyl-Cytosine Dioxygenase TET3 and the Methyl-CpG-binding protein MECP2 Controls Neuronal Maturation"

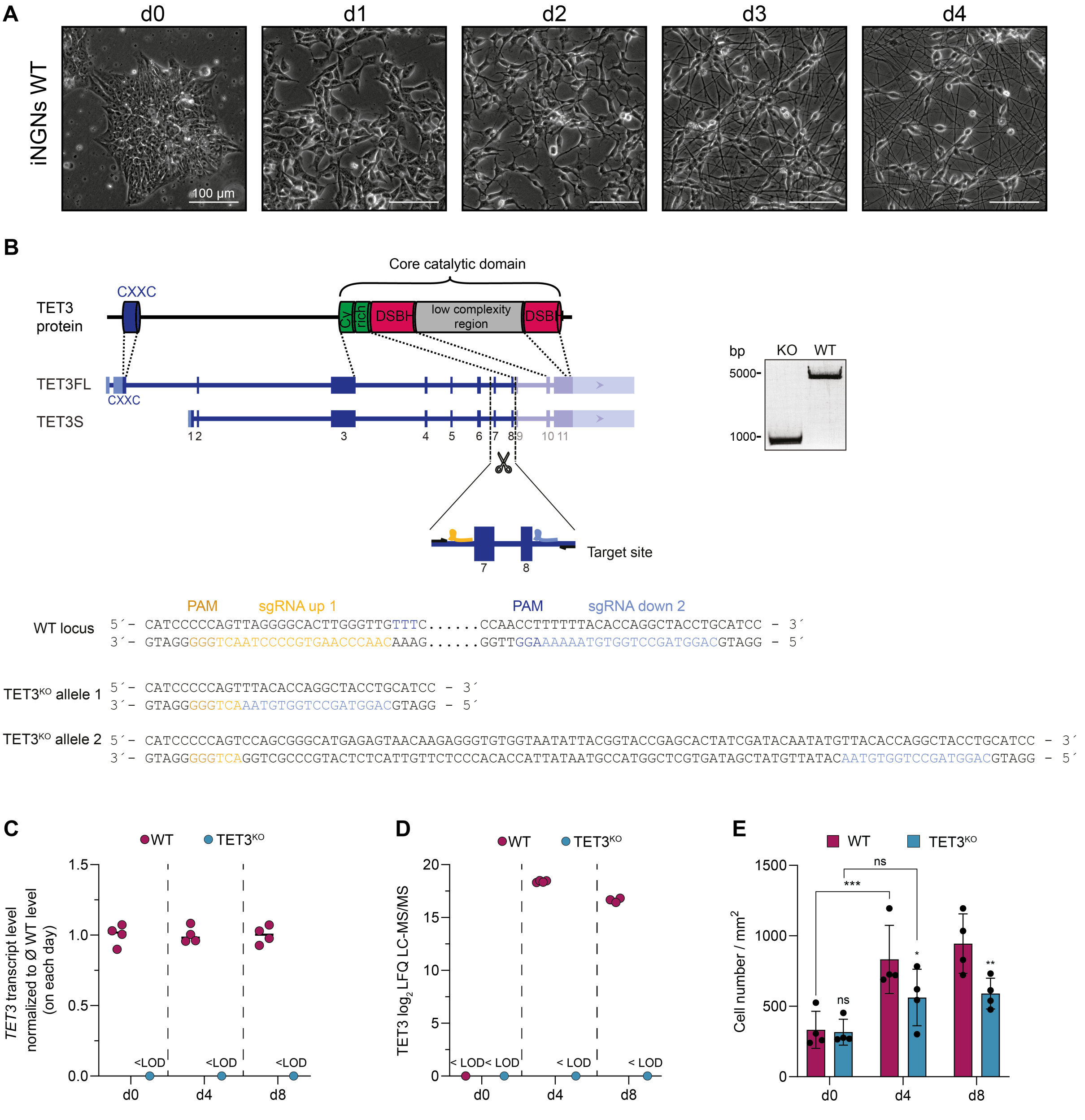


Figure S1: **Characterization of WT and TET3$^{KO}$ iNGNs**. A) Brightfield microscopy images of WT iNGNs before doxycycline induction (d0) and on d1 – d4 after induction. B) CRISPR-Cas9 strategy for generation of TET3$^{KO}$ iNGNs and PCR amplification of the genomic region flanking the CRISPR target site in WT and TET3$^{KO}$ iNGNs. WT alleles result in a 4601 bp band, while the TET3$^{KO}$ allele yields an ~852 bp product. The scheme on the right-hand side depicts the position of the sgRNAs in yellow and blue as well as the primers used in the PCR assay as black arrows. Sanger sequencing of the PCR bands confirmed the deletion of exon 7 and 8 in the TET3$^{KO}$ iNGN cell line resulting in slightly different sequences at the two *TET3* alleles. The target sequence of upstream sgRNA 1 and downstream sgRNA 2 are depicted in yellow and light blue, respectively. The protospacer adjacent motif (PAM) next to each target sequence is colored in darker shades. C) qRT-PCR analysis using primers spanning the CRISPR target site confirms loss of *TET3* transcript in TET3$^{KO}$ cells. Expression was normalized to housekeeping genes and is shown relative to WT. D) Quantification of TET3 protein levels by LC-MS/MS. Peptides mapping to TET3 were detected in WT but were absent in $TET3^{KO}$ samples, indicating the absence of TET3 at the protein level. E) Cell density (cells/mm²) quantified at d0, d4, and d8 of differentiation in WT and $TET3^{KO}$ iNGNs. Bars represent mean ± S.D. Statistical analysis was performed using ordinary two-way ANOVA followed by Tukey's multiple comparisons test (between genotypes at each time point and across time points within genotypes). Details are given in Supplementary Table 1. C), D) LOD = limit of detection. C) – E) Each dot indicates a biologically independent replicate.

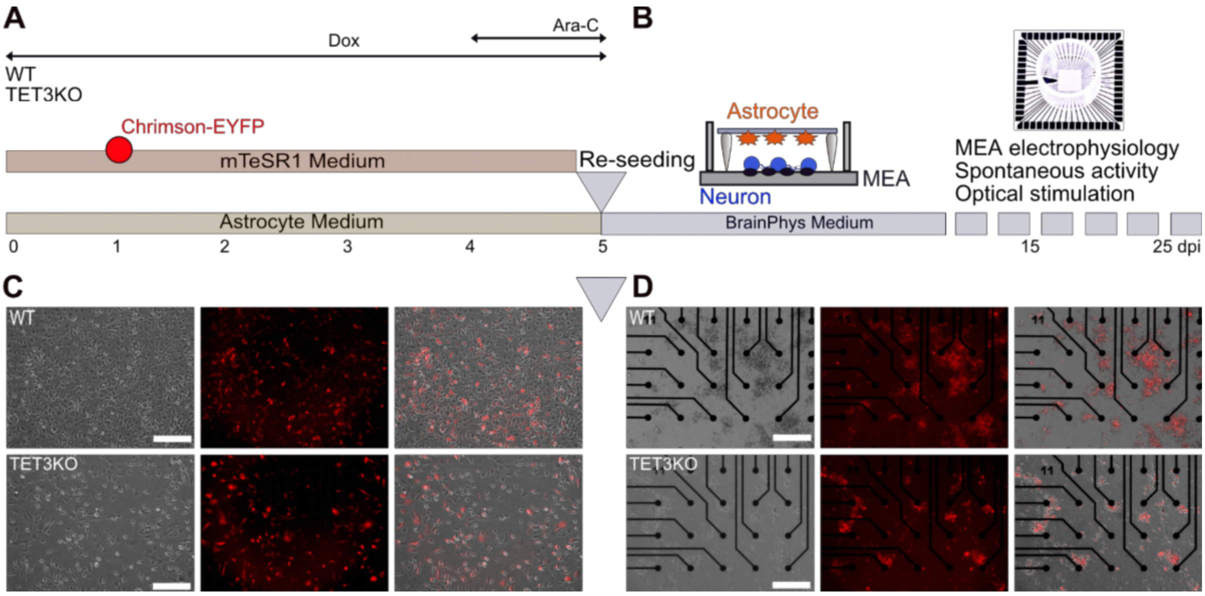


Figure S2: **Re-seeding induced neurons on MEA chips for electrophysiology recordings and optogenetic stimulation.** A) Protocol of neuronal cell induction and optogenetic expression. B) Re-seeding neurons and providing banker culture systems on MEA chips followed by recording spontaneous and optogenetically-evoked activities at d15 and d25. C) Morphology of WT and $TET3^{KO}$ on matrigel coated plates after Ara-C treatment confirmed neuronal phenotype and expression of the ChRimson-EYFP. D) Further differentiation of neurons and development of neuronal networks on MEA chips after re-seeding at d15 with consistent expression of ChRimson EYFP.

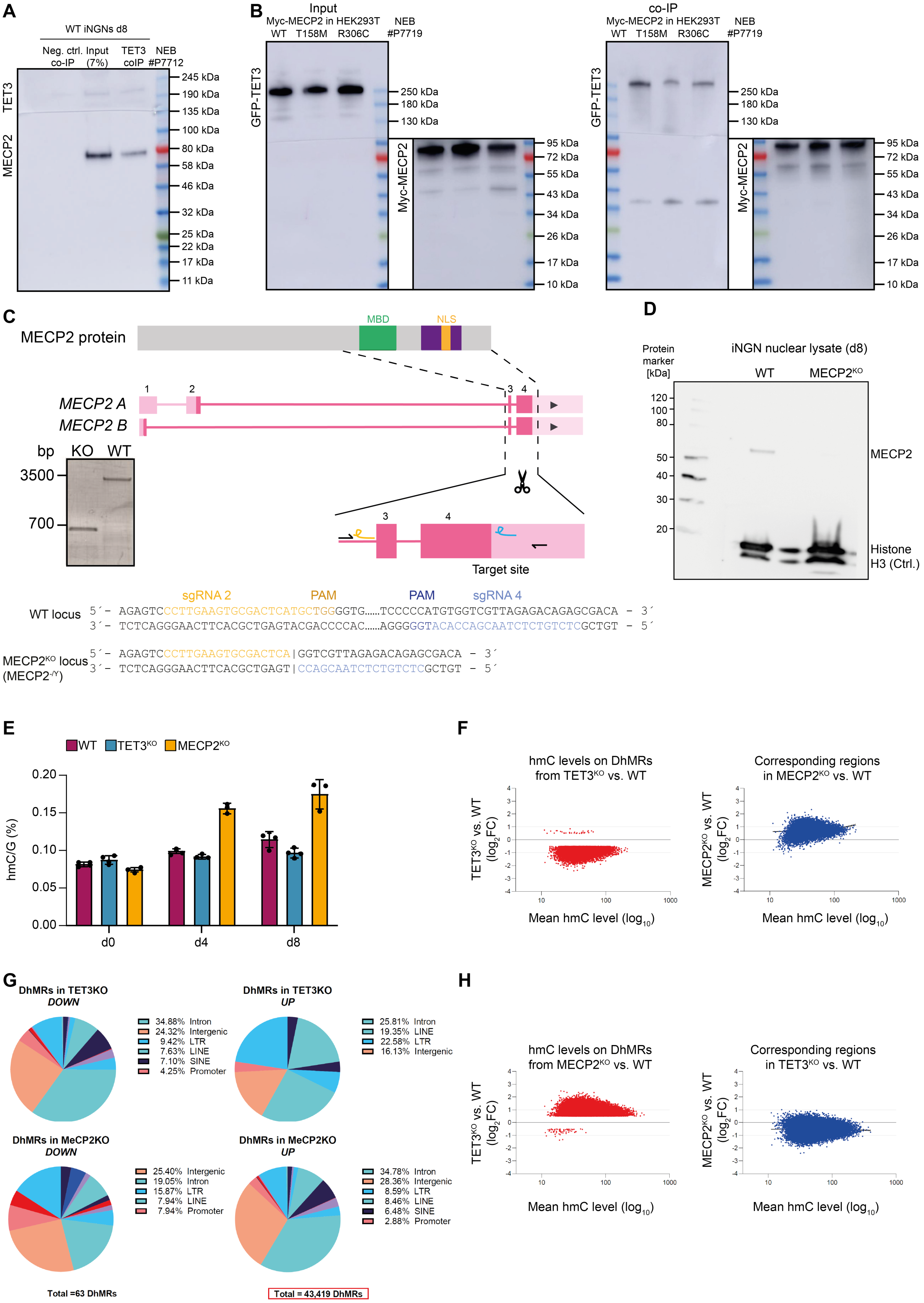

Figure S3: **Functional interplay between TET3 and MECP2**. A) Immunoblot following endogenous TET3 co-immunoprecipitation (coIP) from nuclear lysates of d8 iNGNs. After transfer, the membrane was cut at the 135 kDa marker; the upper part was probed with anti-TET3, and the lower part with anti-MECP2 antibody. B) Immunoblot of Myc-coIP from HEK293T cells transfected with GFP-TET3 and Myc-tagged MECP2 constructs. After blotting, the membrane was cut at the 95 kDa marker; the upper section was probed with anti-GFP, and the lower with anti-Myc antibody. Panels (A) and (B) show the full immunoblot images corresponding to the cropped sections in Fig. 3B and 3C. C) CRISPR-Cas9 strategy for generation of MECP2$^{KO}$ iNGNs. Left: PCR amplification of the genomic region flanking the CRISPR target sites in WT and MECP2$^{KO}$ iNGNs. WT alleles produce a ~3500 bp amplicon; MECP2$^{KO}$ alleles yield a ~700 bp product. Right: schematic representation of sgRNA target sites (yellow and blue) and PCR primers (black arrows). Sanger sequencing confirmed deletion of exons 3 and 4 in the MECP2$^{KO}$ line. D) Immunoblot confirming loss of MECP2 protein in nuclear lysates of d8 MECP2$^{KO}$ iNGNs. Histone H3 served as loading control. E) Levels of hmC/G (%) obtained by QQQ-MS as shown in Fig. 3E but showing the individual values per sample. F) Scatter plot of $log_2$FC of DhMRs (15,506) in d8 TET3$^{KO}$ vs. WT iNGNs and corresponding $log_2$FC in MEP2$^{KO}$ vs. WT. G) Genomic distribution of differentially hydroxymethylated regions (DhMRs) in TET3$^{KO}$ and MECP2$^{KO}$ iNGNs at d8 compared to WT. H) Scatter plot of $log_2$FC of DhMRs (43,482) in d8 MECP2$^{KO}$ vs. WT iNGNs and corresponding $log_2$FC in TET3$^{KO}$ vs. WT.

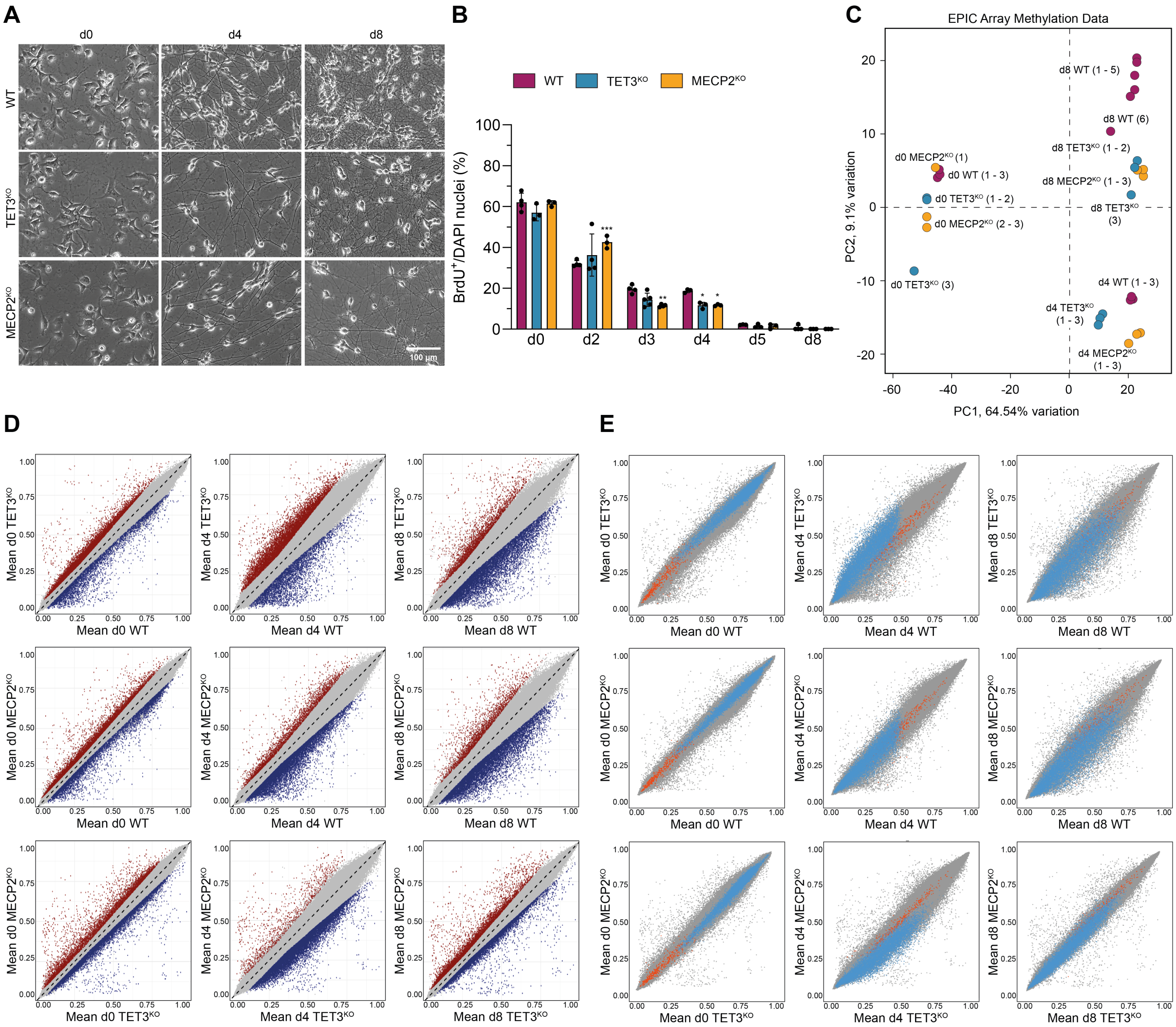


Figure S4: **Phenotypic and DNA methylation profiling of WT, TET3$^{KO}$, and MECP2$^{KO}$ iNGNs during differentiation.** A) Representative brightfield microscopy images of WT, TET3$^{KO}$, and MECP2$^{KO}$ iNGNs at d0, 4, and 8 of differentiation. B) BrdU-incorporation from d0 to d8 across all three genotypes, indicating cell proliferation dynamics. Statistical analysis was performed using ordinary two-way ANOVA followed by Tukey's multiple comparisons test (between genotypes at each time point). Details are given in Supplementary Table 1. C) PCA of genome-wide DNA methylation profiles (EPIC array) for WT, TET3$^{KO}$, and MECP2$^{KO}$ iNGNs at d0, d4, and d8. D) Correlation plots of the top 10,000 differentially methylated CpG sites, comparing methylation patterns between genotypes at d0, d4, and d8. Significantly differentially methylated CpG sites are highlighted in color. E) Correlation plot highlighting methylation dynamics of the top 10,000 CpG sites that undergo significant methylation changes in WT iNGNs between d0 and d4, and how these sites behave in the respective KO lines.

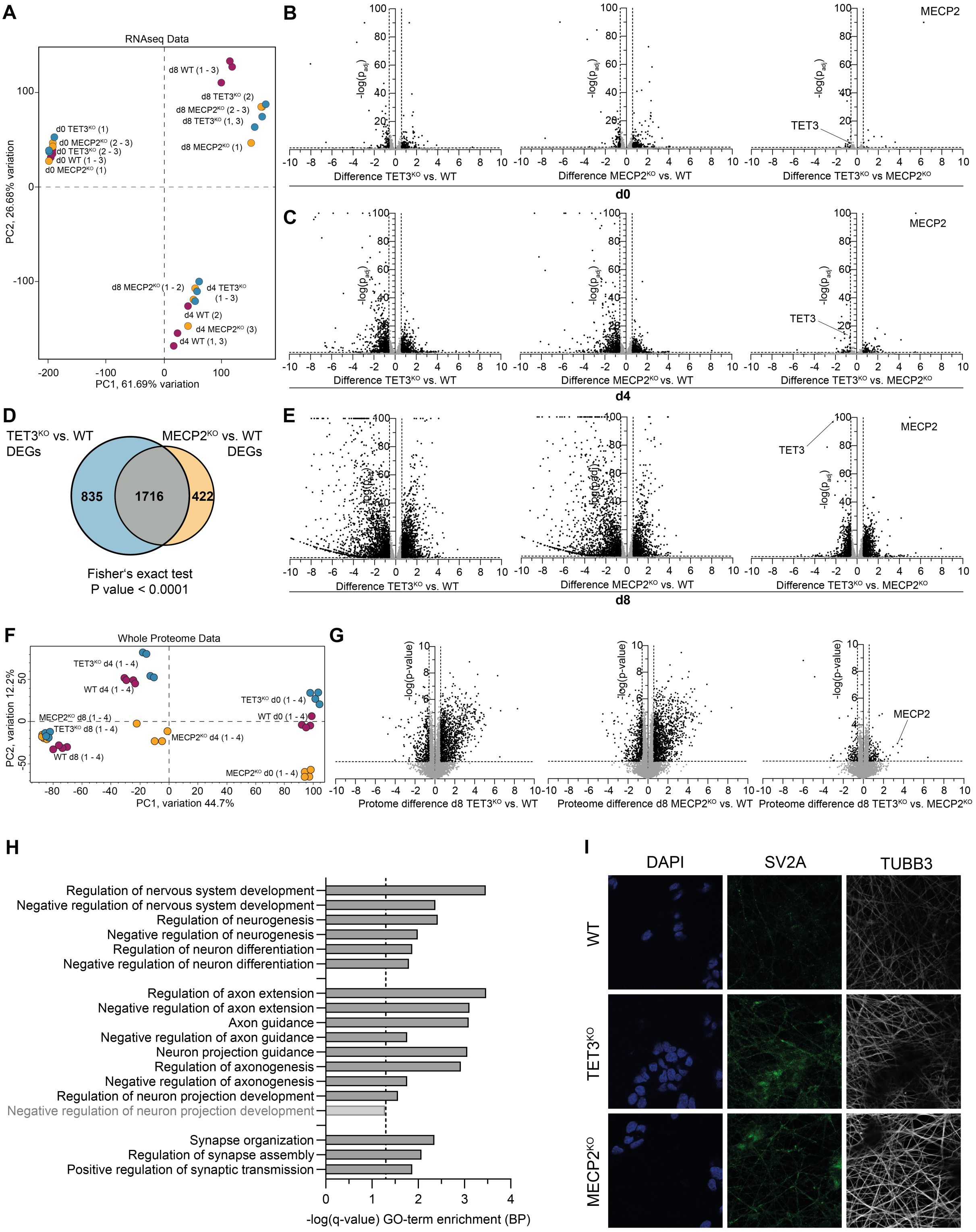


Figure S5: **Transcriptomic and proteomic profiling of WT, TET3$^{KO}$, and MECP2$^{KO}$ iNGNs during neuronal differentiation.** A) PCA of transcriptome profiles (RNA-seq) of WT, TET3$^{KO}$, and MECP2$^{KO}$ iNGNs at d0, d4 and d8 of differentiation. B, C, E, G) Volcano plots showing differential gene or protein expression between indicated genotypes. Panels B, C, and E show RNA-seq results ($log_2$-transformed FPKM values) comparing TET3$^{KO}$ vs. WT, MECP2$^{KO}$ vs. WT, and TET3$^{KO}$ vs. MECP2$^{KO}$ at d0 (B), d4 (C) and d8 (E). Panel G displays proteomics data ($log_2$-transformed LFQ intensities) comparing TET3$^{KO}$ vs. WT, MECP2$^{KO}$ vs. WT and TET3$^{KO}$ vs. MECP2$^{KO}$ at d8. For transcriptomic data, significance was assessed using a two-sided t-test with Benjamini-Hochberg correction. For proteomics, significance was determined by t-test with permutation-based false discovery rate (FDR) estimation. Thresholds for significance in all panels were set to $|log_2FC| > 0.58496$ and $-log(p\text{-value}) > 1.3$. For visualization purposes, $-log(p_{adj})$ values exceeding 100 were capped at 100. Significantly differentially expressed genes or proteins are shown in black; non-significant features are shown in grey. D) Venn diagram of differentially expressed genes (DEGs) at d8 comparing TET3$^{KO}$ and MECP2$^{KO}$ to WT, illustrating the extent of overlap between the two KO conditions. F) PCA of global proteome profiles (DIA, LFQ) of WT, TET3$^{KO}$, and MECP2$^{KO}$ iNGNs at d0, d4, and d8. H) GO term enrichment analysis (Biological Process category) of proteins significantly upregulated in both TET3$^{KO}$ and MECP2$^{KO}$ compared to WT at d8. I) Representative immunofluorescence images used for quantification of synaptic vesicles per neurite length (related to Fig. 4I). SV2A and TUBB3 staining were used to define synaptic vesicles and neurites, respectively.
